# Phagocytic podosomes enable efficient uptake of *Candida auris* by primary human macrophages

**DOI:** 10.1101/2025.09.13.676015

**Authors:** Konstantin Sopelniak, Rawan Batlouni, Qi-fan Sun, Pasquale Cervero, Stefan Linder

## Abstract

The yeast *Candida auris* is an emerging pathogen of steadily increasing importance. Understanding the molecular mechanisms of *C. auris* uptake and intracellular processing by immune cells such as macrophages is thus critical for counteracting the spreading of respective infections. Here, we show that phagocytosis of *C. auris* cells by primary human macrophages involves the formation of dot-like F-actin-rich structures at *C. auris*-containing phagosomes that we characterize as phagocytic podosomes. We analyse the composition, architecture, and dynamics of these structures, showing that they constitute a specific and highly dynamic adaptation of the phagocytic actin network of macrophages. We also show that disruption of phagocytic podosomes is associated with reduced internalization of *C. auris* cells and delayed maturation of respective phagosomes. Our data provide novel insights into the uptake mechanism and cytoskeletal rearrangements upon internalization of *Candida* by immune cells, while also challenging the dogma of a generally uniform and continuous actin network within phagocytic cups. At the same time, we identify *C. auris* as the first pathophysiologically relevant target whose internalization involves the formation of phagocytic podosomes.

## Introduction

The yeast *Candida auris* is a rapidly emerging health threat worldwide ^1^. Since its first isolation from the ear of a Japanese patient in 2009 ^2^*, C. auris* has been found in more than 30 countries on all continents and has accordingly been placed on the list of urgent threats in the 2019 Antimicrobial Threats Report by the Center of Disease Control ^3^. *C. auris* is singular among known *Candida* species, as it can be transferred by humans, enabling patient to patient transmission, which can result in nosocomial outbreaks ^4,5^. Treatment of *C. auris* infections can be difficult, as the fungus shows high resistance against common antimycotics. Accordingly, >90% of known isolates have high resistance rates against fluconazole, while ∼50 of all isolates show high rates of resistance against voriconazole, and ∼10% exhibit also reduced susceptibility against echinocandins ^6^. In contrast to its close relative *C. albicans, C. auris* is mostly present as single cells and tends to form aggregates rather than hyphae ^7^. Moreover, it shows little colonization of the gastrointestinal tract but reaches a higher and more persistent burden on the skin ^8^.

*C. auris* is able to form biofilms, which contributes to its pathogenicity, leading to transmission by infected medical devices ^9^. This is, at least in part, due to its unique form of cell attachment through the recently identified surface colonization factor 1 ^10^. A further property that complicates efficient clearance of infecting *C. auris* cells by the immune system is its unique pattern of cell wall mannosylation, which contributes to its evasion of phagocytosis and efficient killing by human neutrophils ex vivo and in a zebrafish model ^11^. In line with these results, primary human neutrophils show >3x lower phagocytosis rates of *C. auris* cells, compared to uptake of *C. albicans* cells ^12^.

Given the growing importance of *C. auris* as an emerging pathogen, it is crucial to understand how human immune cells interact with and ultimately neutralize infecting *C. auris* cells. Considering the poor phagocytosis rates of *C. auris* by human neutrophils, we therefore investigated the phagocytic uptake and subsequent intracellular processing of *C. auris* by another important arm of the immune system, primary human macrophages.

In general, phagocytosis of particles such as fungal cells proceeds in several steps, which includes i) recognition and surface attachment of the particle, ii) activation of particle internalization, iii) formation of a phagocytic cup, with remodelling into a phagosome upon cup closure, iv) phagosome maturation by fusion with lysosomes, followed by acidification, enabling the activitiy of lytic enzymes, with subsequent degradation of the particle ^13^, and v) resolution of the phagolysosome, enabling the recycling of respective components and lysosome reformation ^14^.

Dynamic remodelling of the actin cytoskeleton is of crucial importance for growth and closure of the phagocytic cup, the cellular protrusion that enables close attachment and eventual engulfment of respective phagocytic particles ^15^. Despite earlier evidence ^16^, this phagocytic actin network has conventionally been seen as largely uniform and continuous. However, recent evidence points to a potential alternative scenario, and several reports have now demonstrated the existence of micron-sized, F-actin-rich dots that resemble “classic”, matrix-associated podosomes at phagocytic cups ^17^. Respectively used cell models include immune cells such as primary human macrophages ^18–20^, murine peritoneal macrophages ^16^, murine bone marrow-derived macrophages ^21^, murine bone marrow-derived dendritic cells ^21^, the murine macrophage cell line RAW 264.7 ^18,19,21,22^, and the human neutrophil cell line HL-60 ^21^ (Table 1).

Of note, all of these experiments were performed using artificial phagocytic particles such as latex beads opsonized with complement ^18^ or IgG ^18,20^, deformable acrylamide beads ^21^, as well as the 2D model of frustrated phagocytosis ^18,19,22^. It is thus currently unclear whether phagocytic podosomes are also formed during the uptake of (patho-)physiologically relevant particles ^17^.

We now show that uptake of *Candida auris* cells by primary human macrophages involves the formation of dot-like F-actin-rich structures that we characterize as phagocytic podosomes. We analyse the composition and architecture of these structures, their dynamics and susceptibility to pharmacological intervention, as well as the consequences of phagocytic podosome disruption on intracellular processing of *C. auris* cells. Our data provide novel insights into the uptake mechanism and cytoskeletal rearrangements upon internalization of *C. auris* by human macrophages. Vice versa, *C. auris* is the first pathophysiologically relevant target whose uptake is shown to involve the formation of phagocytic podosomes, and not uniform actin networks, in phagocytic cups.

## Results

### Phagocytosis of *C. auris* and *C. albicans* involves the formation of F-actin-rich foci at phagosomes

In order to visualize the actin cytoskeleton during uptake of *Candida auris*, primary human macrophages were incubated with *C. auris* cells at a MOI of 50 for 20 min, fixed and stained for F-actin using Alexa488 phalloidin and either DAPI, to stain yeast DNA, or Calcofluor White, which binds β-1,3 and β-1,4 glucans such as chitins in the inner parts of the yeast cell wall ^23^, with subsequent analysis by confocal microscopy. In both cases, we detected prominent dot-like F-actin accumulations at both nascent and fully formed phagosomes containing *C. auris* cells (Fig. 1A-H). In confocal sections, these F-actin dots showed a diameter of mostly 0.5-1.0 µm and were thus similar in size compared to “classic” podosomes at the ventral substrate-contacting side of macrophages. This similar size range was also evident in respective 3D reconstructions of areas of macrophages, although phagocytic F-actin foci were also found to be more patch-like, in contrast to dot-like ventral podosomes (Fig. 1I-K; Suppl. video 1).

**Figure 1.**
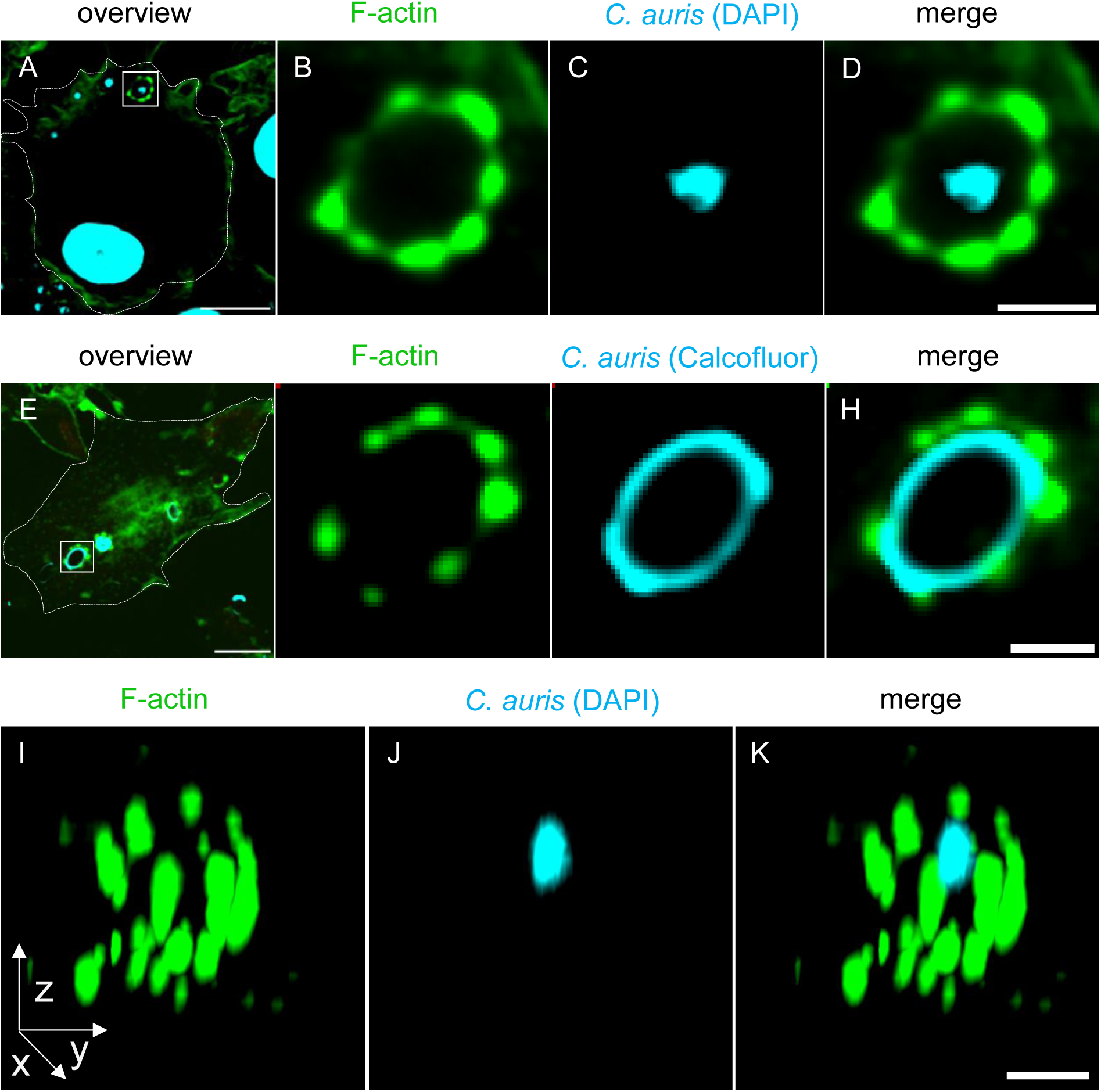
Uptake of *C. auris* by primary macrophages is associated with the formation of micron-sized F-actin foci. (A-H) Confocal micrographs of primary human macrophages infected with *C. auris* cells, stained for F-actin using Alexa488-phalloidin (A,B,E,F) and stained for DNA using DAPI (A,C), or for the *C. auris* cell wall using Calcofluor White (A,G), with respective merges (D,H). White boxes in overview images (A,E) indicate detail regions shown in (B-D, F-H). (I-K) 3D reconstruction of a single *C. auris*-containing phagosome, with F-actin (I), and DAPI staining (J), and merge (K). See also Suppl. Video 1. Scale bars: 10 µm for (A,G). 2 µm for detail images.

To test a potentially more general relevance of F-actin foci during phagocytosis of yeast, we also performed experiments using *Candida albicans* cells. Indeed, also *C. albicans* containing phagosomes showed discrete F-actin foci, which displayed no apparent differences in size, number or distribution to F-actin foci formed at phagosomes containing *C. auris.* However, prolonged incubation of respectively infected cells showed that, in contrast to *C. auris*, phagocytosed *C. albicans* formed hyphae, also within phagosomes, which could lead to eventual break out from phagosomes and disruption of macrophages (Suppl. Fig. 2). The following experiments were thus performed only with *C. auris*.

### *C. auris*-associated F-actin foci share components and architecture with matrix-associated podosomes

Next, we analysed the molecular composition of phagocytic F-actin foci, by staining of endogenous proteins using specific antibodies, and F-actin using Alexa568 phalloidin. In particular, we investigated whether *C. auris*-associated F-actin foci contain components that are typical for ventral podosomes, some of which had also been detected at phagocytic podosomes of phagosomes containing polyacrylamide or latex beads ^17^. We tested components that are typical for all known podosome substructures ^24^, including the F-actin-rich core, the adhesive ring, the regulatory cap structure, the plasma membrane-associated region of the core that is now referred to as the podosome “base” ^25^, and the actomyosin-positive connecting cables thank link individual podosomes (Fig. 2, Suppl. Fig.3). Confocal analysis showed that the podosome core components Arp2 ^26^ and cortactin ^27^ also localized to *C. auris*-associated F-actin dots and showed prominent colocalization with the respective F-actin signal (Fig. 2B1, Suppl. Fig. 2A) The ring components vinculin ^28^ (Fig, 2B2), talin-1 ^28^ (Suppl. Fig. 3B), and αM integrin, a subunit of aMβ2 integrin ^29^ (Fig. 2B3) were also found to be enriched at *C. auris* F*-*actin dots, with local intensity maxima adjacent to the F-actin core, as is typical also for ventral podosomes ^24^. Vinculin is a mechanosensitive protein that is recruited to cryptic binding sites of talin that become exposed upon mechanically stretching ^30^, indicating that phagocytic podosomes could be mechanosensitive. We further detected typical components of the podosome cap, including α-actinin ^31^ (Fig. 2B4), zyxin ^32^ (Suppl. Fig, 3C), and LSP1 ^33^ (Suppl. Fig. 3D). Consistent with their localization at a cap structure, respective fluorescence signals were found to surround the core and extend more towards the cytoplasm on all sides, compared to the actual core. With the unconventional myosin myo1f ^21^ (Fig. 2B5), the Arp2/3-activating nucleation promoting factor WASp ^34^ (Suppl. Fig. 3E), the matrix metalloproteinase MT-1-MMP ^35^ (Fig.2B6), and DNase X ^36^ (Fig. 2B7), we also detected components of the podosome base at respective locations near the plasma membrane). Moreover, we detected prominent staining for myosin IIA between individual F-actin foci, which appears to be consistent with the localization of myosin IIA at the connecting cables of ventral podosomes (Fig. 2B8) ^37^.

**Figure 2.**
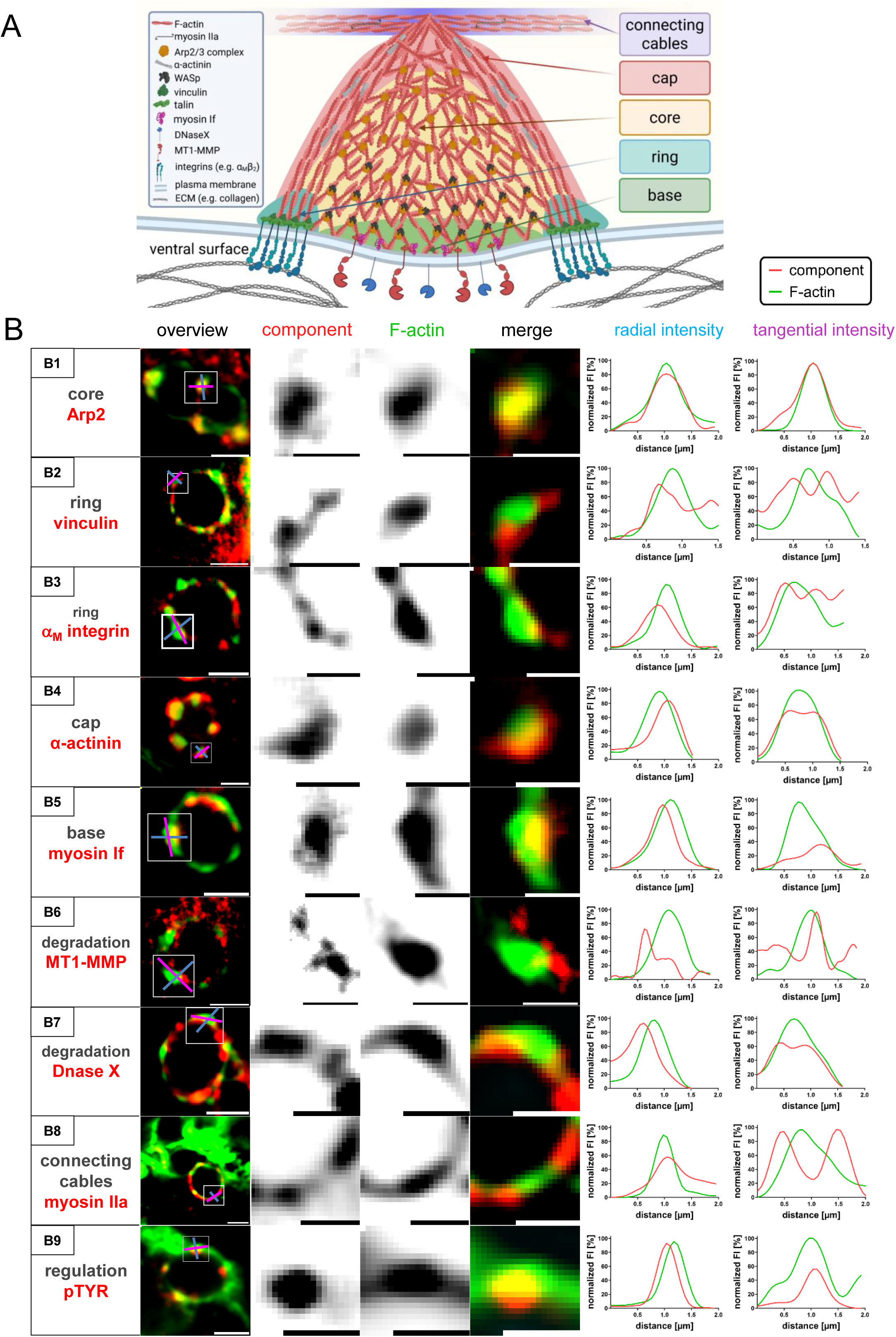
*C. auris-*associated F-actin foci feature components and substructures of ventral podosomes. (A) Podosome model, showing typical substructures, including a cap structure on top of the F-actin core, an adhesion-mediating ring structure, and a base containing plasma membrane-associated proteins. Individual podosomes are linked by connecting cables of actomyosin. Typical components of individual substructures that are found in classic podosomes and were also detected in *C. auris*-associated F-actin foci are listed on the left. Modified from ^24^. (B) Confocal images of *C. auris-*containing phagosomes, with specimens stained for F-actin using Alexa488-phalloidin and for indicated components using specific antibodies. White boxes in overview images indicate detail regions shown on the right, with individual channels in black/white, and colored merges. Graphs show fluorescence intensities gained by respective radial and tangential line scans of individual F-actin foci. From top to bottom: Arp2 (core; B1), vinculin (ring; B2), αM integrin (ring: B3), α-actinin (cap; B4), myosin 1f (base; B5), MT1-MMP (degradative enzyme; B6), DNase X (degradative enzyme; B7), myosin IIA (connecting cables; B8), phospho-tyrosine (regulatory signal; B9); Scale bars: 2 µm for overviews and 1 µm for detail images. See also Suppl. Figure 3.

A further hallmark of ventral podosomes is their regulation by Src tyrosine kinases ^38^ and the resulting enrichment of phosphorylated tyrosine residues in the podosome core structure ^39^. Indeed, use of a phospho-tyrosine specific antibody showed strong accumulation of respective signals at the F-actin structure of *Candida*-associated actin foci (Fig. 2B9; Suppl. Fig. 2F,G). It is noteworthy that these signals were even more sharply defined that the respective F-actin staining. In some cases, they also showed higher relative intensities compared to the respective F-actin signals, which is likely based on the timeline of podosome formation, with Src activity preceding prominent F-actin nucleation ^40^.

Collectively, these experiments showed that F-actin foci associated with *C. auris*-containing phagosomes feature components that are typical for ventral podosomes. Moreover, these components are arranged in the typical podosome architecture, which includes all known podosome substructures such as core, ring, cap, base, and connecting cables. These F-actin foci are also enriched in phospho-tyrosine residues, which are important regulatory signals also of ventral podosomes. In the following, we will thus refer to these structures as “phagocytic podosomes”.

### Formation and dynamics of *C. auris*-associated phagocytic podosomes

We next investigated several parameters of *C. auris-*associated phagocytic podosomes, including the rate of their occurence at phagosomes, their number per phagosome, and their life time. To investigate the time span during the phagocytic process that phagocytic podosomes are present on phagosomes, macrophages were incubated with *C. auris* cells at a MOI of 50 for 5, 10, 20, 40, and 60 min, with subsequent fixation and staining of F-actin by Alexa 568-phalloidin and DAPI to visualize DNA. Phagocytic podosomes were detected at the 5 min time point (Fig. 3A), with 66.4 % ± 4.6 % of phagosomes containing ≥1 podosomes (Fig. 3B), reaching a peak at 10 min, with 75.3 % ± 3.9 % of phagosomes positive for podosomes, and steadily declining at time points 20 min (65.4 % ± 8.7 %) and 40 min (26.0 % ± 9.2 %) (Fig. 3A). 60 min after infection, almost no podosomes were detected at respective phagosomes (4.9 % ± 4.0 % (Fig. 3A). The number of podosomes per phagosome followed a similar pattern, with ∼ 3.0 podosomes per phagosome at time points 5 min, 10 min, and 20 min, and declining at time points 40 min (1.1 ± 0.2) and 60 min (0.2 ± 0.1) (Fig. 3B). Of note, a peak value of 16 podosomes per phagosome was observed at time point 5 min, with further maxima of 10-12 podosomes per phagosome at time points 20 min and 40 min (Fig. 3B), potentially reflecting the maximal numbers of podosomes that can be formed on a *C. auris*-containing phagosome (see Discussion).

**Figure 3.**
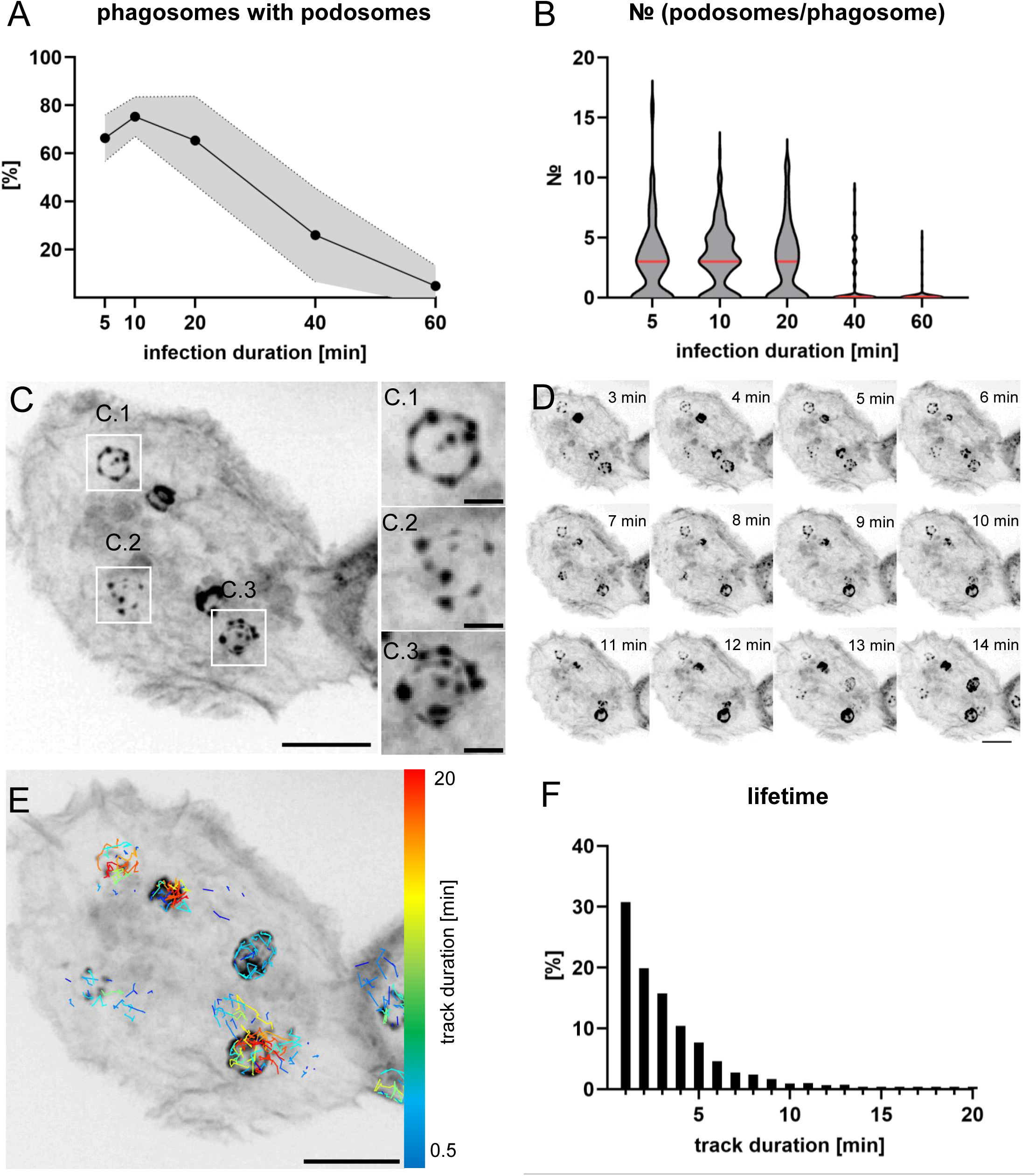
Dynamics of *C. auris*-associated phagocytic podosomes. (A) Graph showing percentage of *C. auris*-containing phagosomes featuring ≥ 1 phagocytic podosome, at the indicated time points post infection. N= 3x30 cells, from each time 3 donors. Values are given as mean ± SEM. (B) Violin plots showing the number of podosomes on individual *C. auris*-containing phagosomes, at the indicated time points post infection. N≥ 3x20 phagosomes, from each time 3 donors. (C-E) Primary macrophage, expressing GFP-Lifeact to visualize F-actin dynamics, internalizing *C. auris* cells; scale bars: 10 µm. (C) Still image from Suppl. Video 2, showing several internalized *C. auris* cells with associated phagocytic podosomes. (D) Gallery of macrophage internalizing *C. auris* cells shown in (C); time after start of experiment is indicated in min:sec. (E) Still image from (C), with tracking of individual phagocytic podosomes; tracks are color coded according to their duration, as indicated by heat bar. (F) Frequency distribution of phagocytic podosome life time at the indicated time points post infection. N= 3x10 cells, from each time 3 donors. Values are given as mean ± SEM. For specific values, see Suppl. Table 1.

In order to visualize the dynamics of phagocytic podosomes, we performed live cell imaging of macrophages expressing GFP-lifeact and infected them with *C. auris* at a MOI of 50. Respective videos confirmed that phagocytic podosomes are already formed at early time points of uptake, when *C. auris* phagosomes are still in the cell periphery of macrophages, and that podosome formation peaks during early phases of phagocytosis (Fig. 3C,D; Suppl. Video 2). Using TrackMate-based analysis, we followed individual phagocytic podosomes from their formation to their dissolution and could thus determine their mean life time as 3.0 ± 2.8 min. A frequency distribution analysis further showed that most (∼60%) of these podosomes had a life time of 1-3 min, with decreasing numbers at further time points. A maximal life span of 18 min was also found for a single podosome (Fig. 3F).

Collectively, these analyses indicate that phagocytic podosomes are formed during early phases of *C. auris* uptake, with the majority of *C. auris-*containing phagosomes showing podosome formation, and that also the number of podosomes per phagosome is highest during early time points of phagocytosis. These data also show that *C. auris*-associated phagocytic podosomes are highly dynamic structures that display fast turnover.

### Phagocytic podosome formation correlates with efficient *C. auris* internalization

The presence of phagocytic podosomes during early phases of *C. auris* internalization by macrophages raised the question of whether these structures support phagocytosis, similar to the “classic” continuous actin network within phagocytic cups ^41^. We thus first tested the impact of various inhibitors that are known to disrupt ventral podosomes, by targeting actin polymerization on several levels, and also by inhibiting Src tyrosine kinase signalling. Tested inhibitors included cytochalasin D, a general inhibitor of actin polymerization ^42^, CK666 and Ck869, two inhibitors of Arp2/3 complex ^43,44^, wiskostatin, an inhibitor of WASp ^45^, as well as PP2, an inhibitor of Src tyrosine kinases and disruption agent for podosomes ^46^. We chose two different inhibitors of Arp2/3 complex based on their slightly diverse activity, as CK666 does not inhibit Arp2/3 complexes containing the ArpC1B subunit, while CK869 does not inhibit Arp2/3 complexes containing Arp3B ^44^. Macrophages were infected with *C. auris* cells at a MOI of 50, incubated for 5, 10, 20, 40, 60, or 80 min, fixed and stained for F-actin and phospho-tyrosine, with ventral podosomes serving as a readout for the efficiency of the treatment (Suppl. Fig. 4)

Respective time courses showed that all five inhibitors effectively disrupted ventral podosomes, in contrast to DMSO controls (Suppl. Fig. 4). At the same time, these treatments led to strong reductions in both the percentage of phagosomes featuring podosomes (Fig. 4A-D) and the number of podosomes per phagosome (Fig. 4E-H). Analysis of *C. auris* internalization showed that cytochalasin D treatment led to a strong decrease in relative internalization (Fig. 4I) and also in the phagocytic index (Fig. 4M), underlining the general importance of actin dynamics in phagocytosis. Inhibition of Arp2/3 complex by CK666 or CK869 led to strong reductions in the assessed parameters. However, neither phagocytic podosome formation (Fig. 4B,F) nor uptake of *C. auris* (Fig. 4J,N) were fully inhibited in the presence of either inhibitor. By contrast, inhibition of WASp, a podosome-localized nucleation promoting factor for Arp2/3 complex ^26,34^ by wiskostatin, led to complete inhibition of phagocytic podosome formation (Fig. 4C,G), and to only residual uptake of *C. auris* cells (Fig.4K,O). As expected, PP2 treatment did not only lead to a strong reduction of ventral podosomes but also to a strong decrease in cellular phospho-tyrosine levels (Suppl. Fig. 3). PP2 treatment led to strongly reduced formation of phagocytic podosomes (Fig. 4D,H) to a delay in internalization (Fig. 4L) and to a ∼50% reduction of the phagocytic index at all measured time points (Fig. 4P), compatible with the impact of Src tyrosine kinases on phagocytosis ^47^ and also on podosome regulation ^48^.

**Figure 4.**
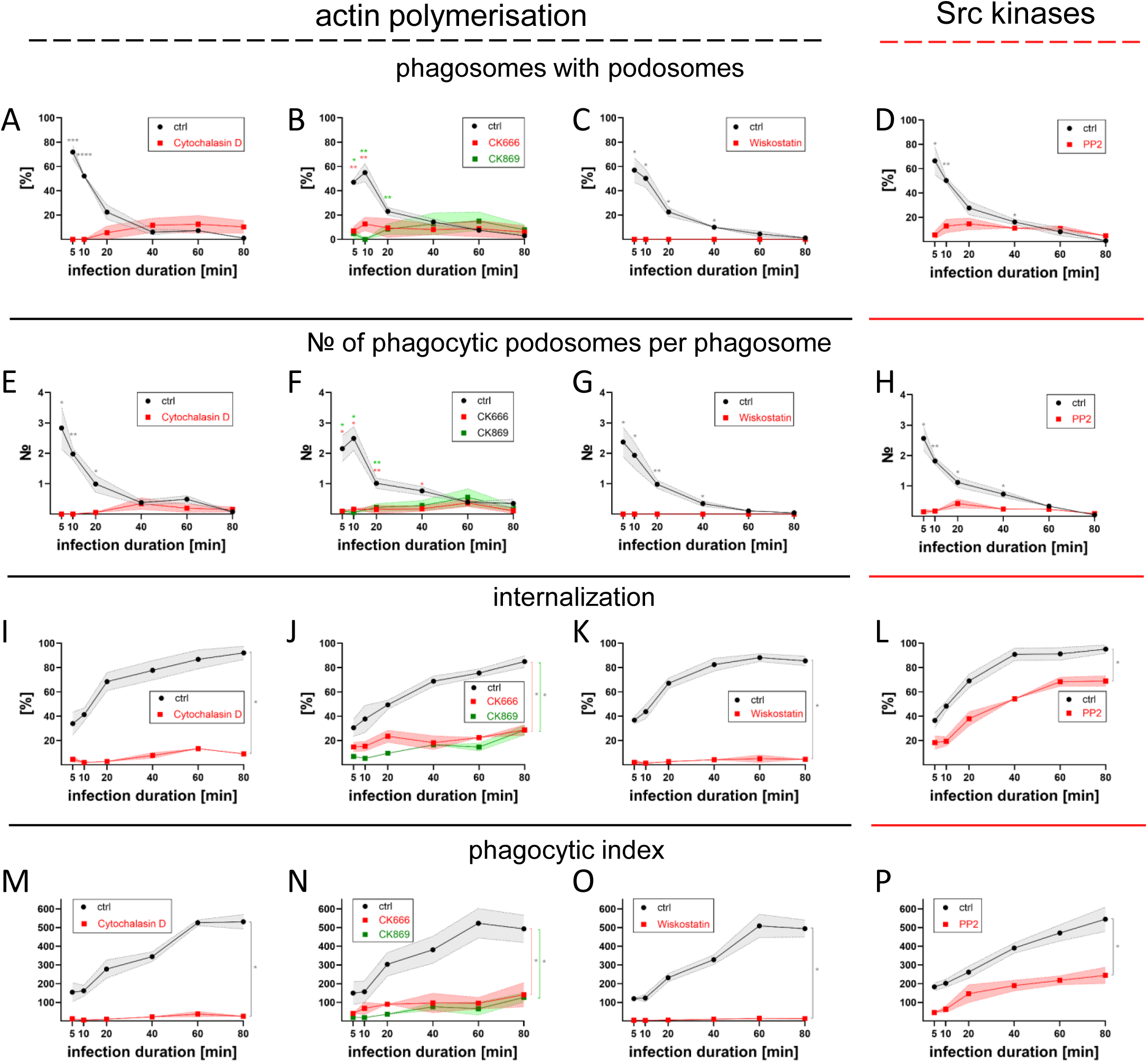
Phagocytic podosomes are regulated by WASp-Arp2/3 complex-based actin nucleation and Src tyrosine kinase activity; disruption of phagocytic podosomes is associated with reduced *C. auris* internalization.. Primary macrophages were treated with cytochalasin D, to inhibit actin dynamics (A,E,I,M), with CK666 or CK869, inhibitors of Arp2/3 complex (B,FJ,N). with wiskostatin, an inhibitor of WASp (C,G,K,O), or with PP2, an inhibitor of Src tyrosine kinases (D,H,L,P). Respective graphs show the percentage of *C. auris*-containing phagosomes featuring ≥ 1 phagocytic podosome (A-D), the number of phagocytic podosomes per phagosome (E-H), internalization rates (%) (I-L), or the phagocytic index (M-P) at the indicated time points post infection. N= 3x30 cells, from each time 3 donors. Values are given as mean ± SEM; *P<0.05; **P<0.01, according to Student’s t-test (A-H), or Wilcoxon test (I-P). For specific values, see Suppl. Table 1.

Collectively, these data show that inhibition of actin polymerization, and in particular of WASp-dependent actin nucleation by Arp2/3 complex, or inhibition of Src tyrosine kinase signalling led to strong reductions in the formation of phagocytic podosomes. This was accompanied by comparable reductions in the uptake of *C. auris* cells by macrophages, indicating a likely involvement of phagocytic podosomes in *C. auris* internalization by macrophages.

### Phagocytic podosomes establish close contact with *C. auris* cells

A main function of podosomes is close adhesion to the underlying surface, through adhesion receptors such as integrins. As *C. auris* associated phagocytic podosomes contain αMβ2 integrin, we tested whether inhibition of phagocytic podosome formation by inhibitors such as CK666 or PP2 or by inhibiting integrin activation by EDTA, through chelation of divalent cations ^18^, resulted in more loosely attached phagocytic cups. For this, we stained the outer region of the *C. auris* cell wall by adding concanavalin A, which binds mannans that are exposed on the yeast surface ^49^, to macrophages ingesting *C. auris* cells. Of note, externally added concanavalin A can only diffuse into incompletely closed cups. This results in a gradient within phagocytic cups, and thus in lower staining intensity at the distal parts of phagocytic cups. We took advantage of this to determine the most basal site of respective cups, which was then used as a reference point to measure the distance between the *C. auris* cell wall and the enclosing actin cytoskeleton. Of note, addition of the various inhibitors resulted in only residual formation of phagocytic cups (see Fig. 3). Associated F-actin stainings in cups were mostly uniform and continuous, thus most likely representing unbranched actin filaments, and not phagocytic podosomes (Fig. 5A-P).

**Figure 5.**
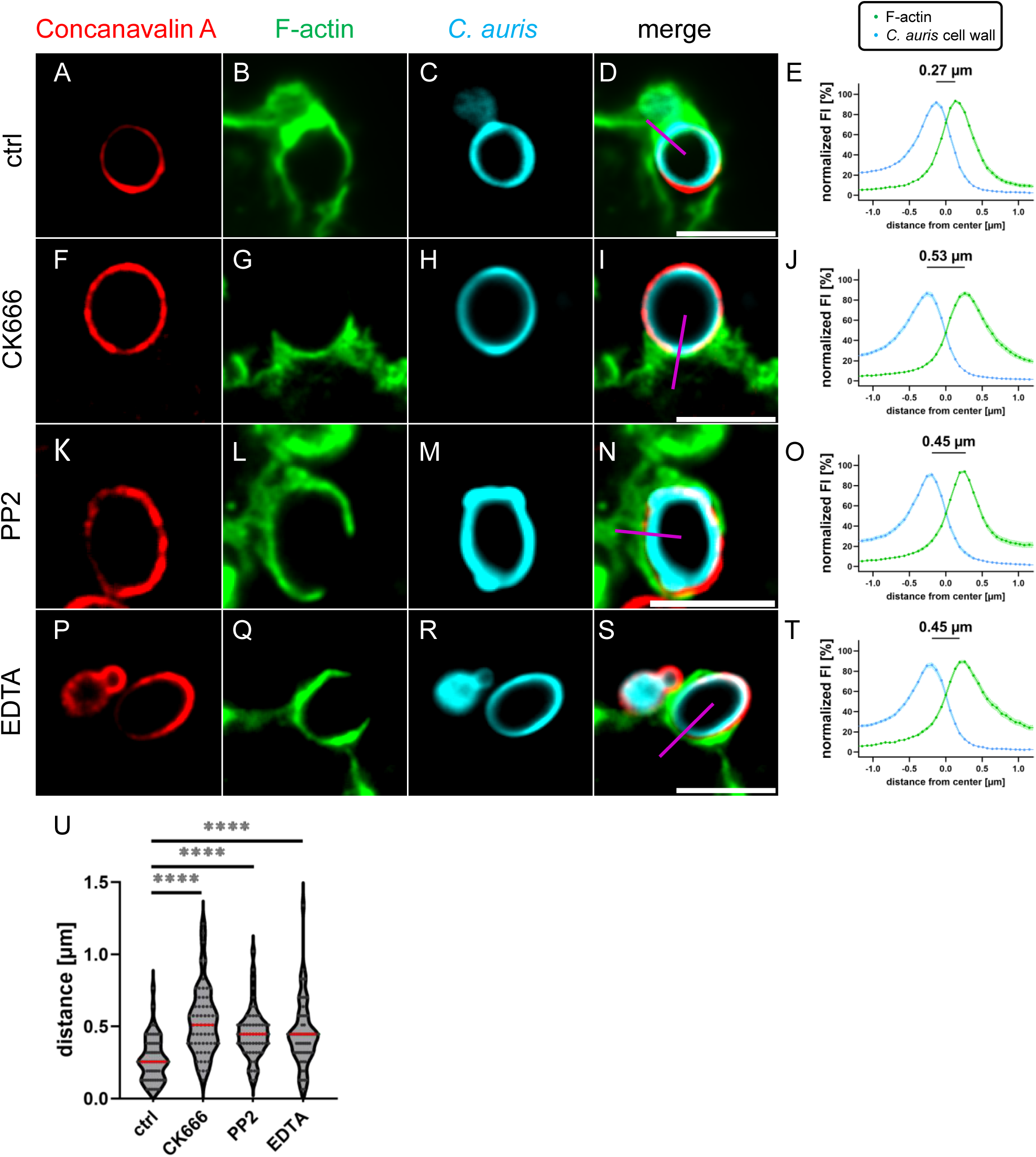
Inhibition of phagocytic podosomes is correlated with compromised attachment of *C. auris* cells to phagocytic cups. Confocal micrographs of primary human macrophages treated with indicated inhibitors and infected with *C. auris.* Specimens are stained for outer and inner cell wall of *C. auris* by Concanavalin A (red; A,F,K,P) or Calcofluor White (blue; C,H,M,R), respectively, and for F-actin using AlexaFluor488-phalloidin (green; B,G,L,Q), with respective merges (D,I,N,S). Cells were treated with CK666, to inhibit Arp2/3 complex (F-I), PP2, to inhibit Src kinases (K-N), or EDTA, to inhibit integrins (P-S). Note that externally added Concanavalin A was used during the experiment to stain only incompletely closed phagocytic cups-The most basal site of phagocytic cups, visible as a faint Concanavalin A staining and indicated by purple lines in merged images, was then used as a reference point to measure the distance between the *C. auris* cell wall and the enclosing actin cytoskeleton. Note also that F-actin stainings in residually formed phagocytic cups are uniform and continuous, thus likely representing unbranched actin filaments, and not phagocytic podosomes. Scale bars: 5 µm. (E,J,O,T) Line scans of fluorescence intensities of the outer cell wall of *C. auris* and the F-actin cytoskeleton of the phagocytic cup, with respective interdistances shown in (U). N= 3x30 cells, from each time 3 donors. Values are given as mean ± SEM. For specific values, see Suppl. Table 1.

In control cells, the mean distance between the *C. auris* cell wall and the underlying actin network was 0.27 µm ± 0.02 µm (Fig. 5A-E). Inhibition of Arp2/3 complex by CK666 resulted in a mean distance of 0.53 µm ± 0.03 µm (Fig. 5F-J), inhibition of Src kinases by PP2 in a mean distance of 0.45 µm ± 0.02 µm (Fig. 5K-O), and inhibition of integrin activation by EDTA resulted in a mean distance of 0.45 µm ± 0.03 µm (Fig. 5P-T), representing relative increases of 96 %, 67 %, and 67 %, respectively (Fig. 5Q). These results show that inhibition of phagocytic podosome formation results in increased distance between the phagocytic actin network and *C. auris* cells as the phagocytic target.

### Disruption of phagocytic podosomes is associated with delayed phagosome maturation

Comparable to their ventral counterparts ^24^, phagocytic podosomes accumulate lytic enzymes such as the matrix metalloproteinase MT1-MMP or DNase X (see Fig. 2), likely through microtubule-dependent transport of respective vesicles. We thus investigated if the presence of phagocytic podosomes influences the maturation of *C. auris* containing phagosomes. For this, macrophages were treated with either CK666 or PP2, infected with *C. auris* cells at a MOI of 50, fixed at time points 5, 10, 20, 40, 60, and 80 min post infection, and stained for LAMP1, to evaluate maturation to phagolysosomes. Compared to controls (Fig. 6A-D), treatment with either CK666 (Fig. 6E-H) or PP2 (Fig. 6I-L) led to reduced LAMP1 recruitment at *C. auris* containing phagosomes at all evaluated time points (Fig. 6 M,N), indicating a delay in phagolysosome maturation.

**Figure 6.**
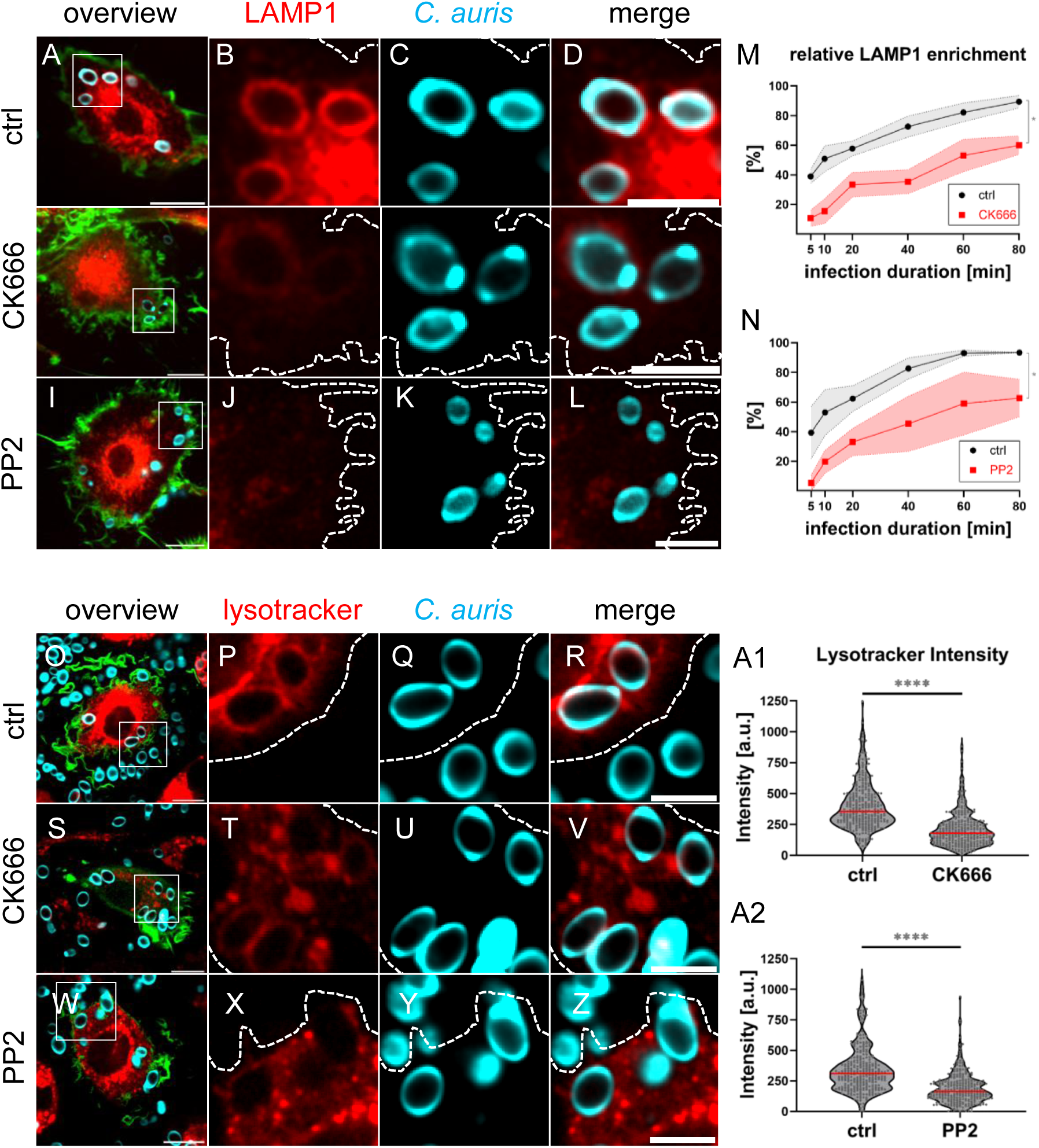
Disruption of phagocytic podosomes is associated with reduced phagosome maturation. (A-L) Confocal micrographs of primary human macrophages treated with indicated inhibitors, infected with Calcofluor White-labeled *C. auris* cells, and stained for LAMP1 using respective antibody at indicated time points post infection. (A,E,I) overview images, with white boxes indicating ROIs shown on the side, including LAMP1 staining (red; B,F,J), *C. auris* (blue; C,G,K), with respective merges (D,H,L). Cells were treated with CK666, to inhibit Arp2/3 complex (E-H), or PP2, to inhibit Src kinases (I-L). (M,N) LAMP1 enrichment at *C. auris* phagosomes at indicated time points post infection, with respective LAMP1-based fluorescence intensity at time point 80 min in control cells set to 100%. N= 30 cells, from each time 3 donors. (O-Z) Confocal still images from live cell videos of primary human macrophages, treated with indicated inhibitors, infected with Calcofluor White-labeled *C. auris* cells, and incubated with lyostracker for detection of acidic compartments. (O,S,W) overview images, with white boxes indicating ROIs shown on the side, including lysotracker staining (red; P,T,X), *C. auris* (blue; Q,U,Y), with respective merges (R,V,Z). Cells were treated with CK666, to inhibit Arp2/3 complex (S-V), or PP2, to inhibit Src kinases (W-Z). Scale bars: 10 µm for overview images, 5 µm for detail images. (A1,A2) Lysotracker enrichment at *C. auris* phagosomes, with respective lysotracker-based fluorescence intensity at time point 1h in control cells set to 100%. N= 3x100 phagosomes, from each time 3 donors. Values are given as mean ± SEM; *P<0.05; ****P<0.001, according to Wilcoxon test (M,N), or Student’s t-test (A1,A2). For specific values, see Suppl. Table 1.

To investigate this further, we next asked if the decrease in LAMP1 recruitment to *C. auris* phagosomes was also accompanied by reduced phagosomal acidification. For this, living macrophages were treated with either CK666 or PP2, infected with *C. auris* cells at a MOI of 50, with addition of lysotracker, a pH-sensitive reporter of acidification. Images of respective cells were recorded at 1h post infection, and lysotracker-based fluorescence intensities around *C. auris* phagosomes were measured. Compared to controls (Fig. 6O-R), treatment with either CK666 (Fig. 6S-V) or PP2 (Fig. 6W-Z) led to ∼ 55% reductions in LAMP1 recruitment at *C. auris* containing phagosomes (Fig. 6A1, 6A2). Collectively, these data show that disruption of phagocytic podosomes at *C. auris* containing phagosomes is associated with a delay in phagolysosome maturation.

## Discussion

In this study, we show that phagocytic uptake of *Candida auris* by primary human macrophages involves the formation of discrete F-actin rich structures with podosomal characteristics. This is the first report to identify phagocytic podosomes in the uptake of fungal cells by human immune cells. Moreover, as previous studies used artificial particles such as latex or polyacrylamide beads ^18,20,21^, we now identify *Candida auris*. as the first pathophysiologically relevant target whose uptake is mediated by phagocytic podosomes.

Our initial experiments showed that F-actin foci are also formed during uptake of *C. albicans* cells, pointing to a more general importance of phagocytic podosomes in the uptake of yeast cells. This seems particularly interesting in light of the reported differences in composition of the *C. auris* versus the *C. albicans* cell wall ^11^. Still, observation of the subsequent fate of internalized yeast cells showed that, in contrast to *C. auris* cells, which remain within macrophage phagosomes or are even degraded, *C. albicans* cells can germinate within phagosomes, break out of macrophages and establish an elaborate hyphal network (Suppl. Fig. 2), as also reported earlier ^50^. Of note, a recent report showed that *C. auris* cells internalized by murine bone marrow-derived macrophages can kill these immune cells by triggering host metabolic stress ^51^, indicating that the ultimate outcome of a respective infection likely also depends on the host species.

Using immunofluorescence, we investigated the molecular composition and substructures of phagocytic podosomes. We found typical components of ventral podosomes, which also localized to respective substructures, including proteins of the cap (α-actinin, zyxin, LSP1), core (Arp2, cortactin), ring (vinculin, talin-1, αM integrin), and base (myo1f, WASp, MT1-MMP, DNase X) ^24^. This inventory shows that phagocytic podosomes share a similar set of components, arranged in a comparable architecture, with their ventral counterparts (Figs. 2,7). Moreover, the connection through actomyosin filaments, similar to the connecting cables of ventral podosomes, indicates that also phagocytic podosomes are part of a higher-ordered array. The presence of Arp2 and WASp further points to the existence of a branched actin network at the podosomes core. This network is likely anchored to the plasma membrane by the phospholipid-binding domain of myosin 1f ^52^. The detection of adhesion plaque proteins and integrin is also consistent with a linkage to the surface of the ingested particle, i.e. *C. auris* cells, while strong signals for phospho-tyrosine indicate that, like classic podosomes, also phagocytic podosomes are regulated by tyrosine kinase signalling (see below).

**Figure 7.**
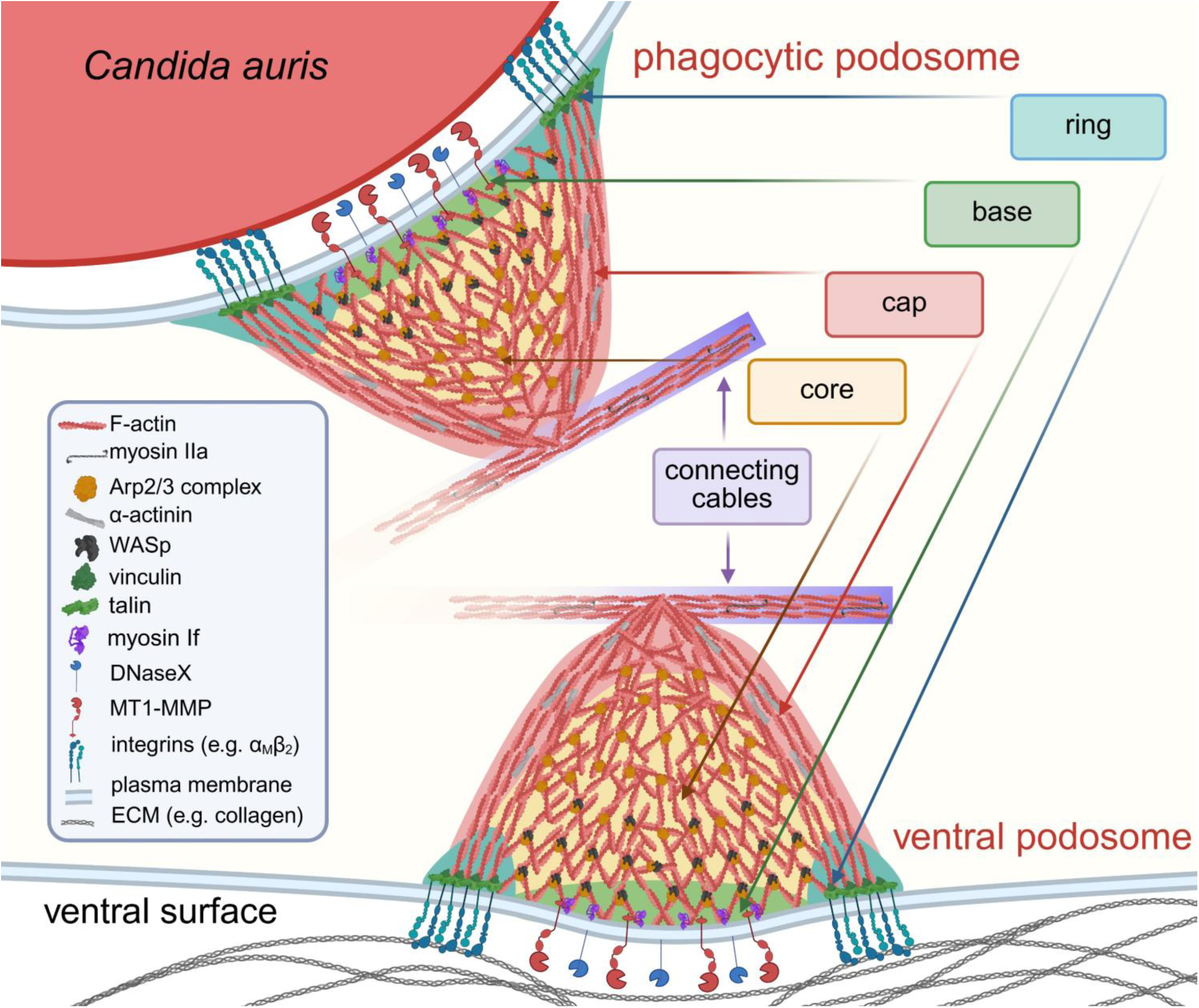
Model: *C. auris*-associated phagocytic podosomes share components and substructures with their “classic”, ECM-associated counterparts. Detected substructures and components include an F-actin-rich core (positive for Arp2 and cortactin), a cap structure on top of the core (positive for α-actinin, zyxin, LSP1), a ring structure mediating adhesion to the phagocytic particle (containing vinculin, talin-1, αµ integrin), and a base near the plasma membrane (containing myosin 1f, WASp, MT1-MMP, DNAse X, and enriched in phospho-tyrosine residues). Individual phagocytic podosomes are linked by actomyosin filaments, similar to the connecting cables of ventral podosomes, thus establishing a flexible podosome array on phagosomes. The high turnover rate of phagocytic podosomes likely enables dynamic restructuring of the nascent phagocytic cup. In combination with establishing close adhesion to the pathogen surface, this likely mediates efficient internalization of *C. auris* cells in macrophages. Recruitment of degradative enzymes such as MT1-MMP or DNase X could, directly or indirectly, contribute to phagosome maturation and particle degradation.

Further analysis showed that *C. auris-*associated podosomes are formed during the early phases of phagocytosis (5-20 min). During this time, the majority of phagosomes exhibit podosome formation, with the highest detected numbers of podosomes per phagosome. Moreover, *C. auris*-associated podosomes were found to be highly dynamic, with a mean life time of 3.0 ± 2.8 min, which is lower than that observed for podosomes on phagosomes containing latex beads (5.0 ± 3.3 min) ^20^, and also lower than the life time of ventral podosomes (9.9 ± 0.6 min) ^33^, with all values measured in primary human macrophages. As the majority of phagosomes feature podosomes during the first 20 min of phagocytosis, this implies a constant turnover of podosomes on individual phagosomes, which could also be seen in respective live cell videos. It is thus likely that any potential function(s) exerted by these podosomes should be associated with early time points of phagocytosis and that they necessitate high adaptability of these structures in time and space.

Considering the potential involvement of phagocytic podosomes in the uptake of target particles, the respective phagosome surface should offer sufficient area and also appropriate stiffness to allow for podosome formation. Therefore, the parameters i) size and ii) rigidity appear to be of particular importance:

1. size: *Candida* cells show a diameter of ∼4 µm. Their surface area, in first approximation calculated for a sphere (π x d^2^), is thus ∼50 µm^2^. *C. auris*-associated phagocytic podosomes show a diameter of ∼1 µm, which is comparable to that of ventral podosomes of macrophages ^24^, which also show a typical spacing of 1.17 µm ^53^. A single ventral podosome thus occupies an area with a diameter of ∼2.17 µm. The surface area covered by each podosome (π x r^2^) thus corresponds to 3.7 µm^2^. Combining these values, the surface of a *Candida* containing phagosome would theoretically enable the formation of 50/3.7 = 13.5 podosomes. Indeed, we found a maximum value of 16 podosomes per phagosome at the time point 5 min after start of phagocytosis, with respective peak values of 10-12 at later time points (Fig. 3B), which is in good agreement with these calculations. This also indicates that podosomes on *Candida* phagosomes can show a similar spacing compared to ventral podosomes. Furthermore, the mean value of podosome number per phagosome was ∼3 during the podosome-forming phase of phagocytosis, indicating that, for most phagosomes, not all of the available surface was covered by podosomes. However, it should also be considered that phagocytic podosomes are not necessarily round, but can also be more patch-like (Fig. 1I-K, Suppl. Video 1). Therefore, phagocytic podosomes could cover a larger surface than an equivalent number of their ventral counterparts.
2. rigidity: Atomic force measurements show that *Candida albicans* cells exhibit stiffness regimes in the range between 183 kPa and 364 kPa, depending on their cultivation on glass or polished titanium ^54^. In comparison, THP-1 monocytic cells have been shown to form podosomes on stiff polyacrylamide (PAA) gels of 323 kPa, but not on softer PAA surfaces of 88 kPa ^55^. The rigidity of *Candida* cells thus seems to be in the range that is required for efficient formation of ventral podosomes. Interestingly, structures resembling phagocytic podosomes in RAW 264.7 murine macrophages ingesting IgG-coated deformable acrylamide beads were formed in greater numbers on softer (1.4 kPa) compared to stiffer beads (6.5 kPA) ^21^. It is currently unclear to which degree differences in species, cell type or primary versus transformed cells impact on the formation of phagocytic podosomes and the required parameters.

During “classic” phagocytosis, the continuous actin network in phagocytic cups drives membrane protrusion, while enabling the local concentration of phagocytic receptors such as integrins that mediate close adhesion to the engulfed particle ^56^, ultimately leading to phagocytic cup closure and internalization of the target ^57^. In this context, it has to be considered that a differently structured phagocytic actin cytoskeleton, in the form of phagocytic podosomes, should fulfill similar functions. Indeed, ventral podosomes exhibit a variety of functions that even exceed those of “classic” phagocytic actin networks, including close adhesion to the substratum, mechanosensing of substrate rigidity and topology, as well as degradation of proteins and DNA ^24^. We therefore investigated potential functions of phagocytic podosomes during uptake of *C. auris* cells, including i) close adhesion to the phagocytic target, ii) a role in the internalization of *C. auris*, iii) mechanosensing of the phagocytic particle, iv) a role in maturation of phagosomes.

1. close adhesion: We detect the integrin α_M_β_2_ in the ring structure of phagocytic podosomes, which points to an adhesive function of phagocytic podosomes, similar to the role of this integrin in matrix-associated podosomes ^58^. Interestingly, α_M_β_2_ has been implicated in the recognition of *C. albicans*, by binding to the mannoprotein Pra1 ^59^, which appears to be in line with the general importance of β2 integrins in phagocytosis of bacterial or fungal pathogens by immune cells ^60^. It is currently unclear whether *C. auris* also expresses Pra1, although a BLAST search (https://www.ncbi.nlm.nih.gov/) indicated the existence of a possible homolog (CJI96_0000822) ^61^. It is also possible that further proteins of the *C. auris* cell wall are recognized by α_M_β_2._ Our measurements of the distance between the *C. auris* cell wall and the actin cytoskeleton within phagocytic cups are in line with the presence of integrin in phagocytic podosomes. Inhibition of phagocytic podosome formation by CK666 or PP2, or inhibition of integrin activation by EDTA led to significantly enhanced distances between the *C. auris* cell surface at the underlying actin cytoskeleton (Fig. 5). This indicates that *C. auris* cells as phagocytic targets are likely more loosely attached to the phagocytosing macrophage. Podosomes within phagosomes apparently mediate closer attachment to the phagocytic target than podosome-free phagosomes (see Fig. 4).
2. internalization: Using various inhibitors of actin polymerization or Src tyrosine kinase activity, we observed significant reductions in the number of podosome-positive phagosomes, the number of podosomes per phagosome, as well as in uptake of *C. auris* cells by macrophages. Upon inhibition of Src tyrosine kinases by PP2, phagocytic podosomes were still formed, although at greatly reduced levels (Fig, 4D,H). This was accompanied by a ∼50% decrease of the phagocytic index (Fig. 4P). Furthermore, inhibition of WASp by wiskostatin resulted in complete disruption of phagocytic podosomes (Fig, 4C,G) and also in almost complete abrogation of internalization (Fig, 4K,O), indicating the importance of WASp-based activation of Arp2/3 complex for *C. auris* internalization. As WASp is localized exclusively to podosomes ^34^ and phagocytic cups ^62^ in macrophages, this result points to a critical role of phagocytic podosomes in *C. auris* uptake. Interestingly, neither inhibitor of Arp2/3 complex, CK666 or CK869, fully disrupted phagocytic podosome formation (Fig. 4B,F) or *C. auris* internalization (Fig. 4J,N). These results could be based on the fact that not all Arp2/3 complexes are targeted by either inhibitor (see above) and could thus indicate the presence of both ArpC1B- and Arp3B-containing complexes at phagocytic podosomes. Still, disruption of phagocytic podosomes by either inhibitor was more efficient than the previously reported partial disruption of phagocytic podosomes during uptake of IgG-coated latex beads, even at high levels of CK666 (150 µm) ^20^. As we used the same cell type in the present study, primary human macrophages, this difference could be due to the different phagocytic targets, latex beads versus *C. auris* cells. Collectively, in all inhibitor experiments, we observed a close correlation between the levels of phagocytic podosomes and the efficiency of *C. auris* uptake, pointing to a central role of phagocytic podosomes in the uptake of *C. auris* cells. Still, it has to be considered that inhibitors such as cytochalasin D or CK666 are not podosome-specific, and alternative pathways of internalization that are independent of phagocytic podosome formation, may also play a role in engulfment of the yeast cells, although this contribution appears to be minor.
3. mechanosensing: Phagocytic podosomes contain contractile and mechanosensitive elements, such as talin-1 or actomyosin cables, that are arranged in a typical podosome architecture of cap, core, ring, and base, which enables mechanosensitive probing of the underlying substratum ^63^. Moreover, the stiffness regime of *Candida* cells lies in the range between 183 kPa and 364 kPa ^54^, similar to the stiffness required for efficient formation of podosomes (323 kPa) ^55^ (see above). The presence of vinculin at phagocytic podosomes further points to the mechanical stretching of its binding partner talin, leading to the exposure of cryptic binding sites for vinculin ^30^. It is also noteworthy that, upon internalization of budding *C. auris* cells, vinculin-rich phagocytic podosomes frequently formed at the neck between mother and daughter cells (not shown), pointing to a potential ability of these structures to detect discontinuities in surface topology, which is also a main function of ventral podosomes ^64,65^. Interestingly, an earlier study showed that stiffer polystyrene beads were ingested more efficiently by phorbol-ester activated RAW 264.7 cells, through integrins being coupled via talin and vinculin to the actin cytoskeleton ^66^. Considering that *Candida* cells are relatively stiff ^54^, and that their efficient uptake is associated with the formation of phagocytic podosomes, which contain integrins, talin, and vinculin, our results seem to be in good agreement with these earlier findings.
4. phagosome maturation: We show that depletion of phagocytic podosomes by CK666 or PP2 inhibitors is accompanied by significantly reduced acquisition of LAMP1, a marker of phagolysosome fusion, and also by reduced acidification of *C. auris* containing phagolysosomes. These findings point to a potential role of phagocytic podosomes also in the maturation of phagolysosomes. It could be envisioned that phagocytic podosomes serve as docking sites for vesicles carrying respective degradative enzymes or parts of the machinery for phagolysosomal maturation. Considering that ventral podosomes are repeatedly contacted by microtubule plus ends ^35,67^, and that degradative enzymes such as MT1-MMP persist at the plasma membrane beyond the life time of individual podosomes ^68^, this should be a line of investigation worth following up in future studies. Still, we cannot exclude that especially inhibition of Src tyrosine kinases by PP2 also works through other, non-podosomal pathways, as Src kinases can also influence the interaction of phagosomes with lysosomes ^69^.

In conclusion, we identify the yeast *Candida auris* as the first pathophysiologically relevant target whose uptake into primary macrophages involves the formation of phagocytic podosomes. At the same time, our results give detailed insights into the uptake mechanisms of *C. auris* cells by primary human macrophages, which should be of particular importance considering that *C. auris* constitutes an emerging health threat. We show that phagocytic podosomes are highly dynamic structures that likely play a crucial role in internalization of *C. auris* cells, by providing close adhesion within the phagocytic cup and enabling efficient uptake. Moreover, the presence of lytic enzymes at phagocytic podosomes points to their potential role in the initiation of phagosome maturation. As classic podosomes exhibit further functions, such as substrate degradation or topography sensing, it will be interesting to see which further roles phagocytic podosomes can play during the phagocytic process.

At the same time, our results contribute to challenging the decades-long dogma of a uniform and continuous actin network in the phagocytic cup. Accordingly, we propose that phagocytic podosomes are not only involved in the uptake of *C. auris* cells, but are likely also important for internalization of phagocytic targets that exhibit similar size and rigidity. Indeed, formation of phagocytic podosomes could be seen as an adaptation of the cytoskeletal machinery of immune cells that is triggered by ingestion of comparably large and rigid particles, ensuring efficient uptake and internal processing of these difficult targets.

## Methods

### *Candida spp* growth conditions

*C. auris* (strain CDC B11903) and *C. albicans* (strain ATCC 10231) were cultivated in LB medium overnight, plated on YPD agar and cultivated at 33 °C overnight. Thereafter, fresh agar plates were sealed with parafilm and kept at 4 °C for up to 2 weeks.

### Primary macrophage cell culture

Primary human monocytes were isolated from buffy coats (kindly provided by Frank Bentzien, Transfusion Medicine, UKE, Hamburg, Germany). 20 ml blood were slowly transferred on 15 ml Lymphocyte Separation Medium 1077 (PromoCell, Heidelberg, Germany) and centrifuged for 30 min at 4 °C and 430 ×g without acceleration and deceleration. Buffy coat layer was transferred in a new falcon and filled up to 50 ml with cold RPMI (Gibco, Paisley, UK). Leucocyte fractions were washed twice in RPMI and centrifuged for 10 min at 4 °C and 430 ×g. Enriched leukocytes were resuspended in 400 μl monocyte buffer (5 mM EDTA and 0.5 % human serum albumin in Dulbecco’s PBS [DPBS], pH 7.4), mixed with 200 μl of magnetic CD14 positive beads suspension (Miltenyi Biotec, Bergisch Gladbach, Germany) and incubated for 15 min on ice. Thereafter, the mixture was loaded on Separation columns LS (Miltenyi Biotec, Bergisch Gladbach, Germany), previously placed in a magnetic holder, and equilibrated with 500 μl cold monocyte buffer. Trapped CD14+ monocytes were washed on column with 1500 μl monocyte buffer and then eluted with 1ml monocyte buffer into 15 ml cold RPMI after removal from the magnets. Monocytes were resuspended in 40 ml RPMI and seeded on a 6-well plate (Sarstedt, Nuembrech, Germany) at a density of 2x10^6^ cells per well. After adhesion of monocytes (2-4 h), RPMI medium was replaced by 2 ml monocyte culture medium (RPMI substituted with 20% human serum and 1% penicillin/streptavidin (Sigma-Aldrich, Missouri, USA)). Monocytes were cultivated in an incubator at 37 °C, 5% CO_2_, and 90% humidity. Isolated monocytes were differentiated for at least 6 days.

### Phagocytosis assays

6- to 14-day old macrophages were detached by incubation with Accutase (Invitrogen, Massachusetts, USA) for at least 30 min in culturing conditions and resuspended in monocyte medium. Macrophages were seeded at a density of 1x10^5^ cells per glass coverslip (12-mm Ø) and incubated for at least 60 min at culturing conditions for at least 80% confluence. Monolayers of macrophages were infected with *C. auris* or *C. albicans* by sedimenting the yeast cells at 200 xg for 3 min. Infection was performed with MOI 50:1 in RPMI at 37 °C, 5% CO_2_, and 90% humidity for 5, 10, 20, 40, 60 min (Fig.3 A,B) or up to 3-24 h (Suppl.Fig.1 A-R) before fixation for 10 min in 3.7% polyformaldehyde (PFA) in PBS and subsequent immunofluorescence staining.

### Use of inhibitors

Macrophages seeded on glass coverslips (12 mm Ø) were starved for 1 h with RPMI 1640 under culturing conditions, preconditioned for 10 min with respective inhibitors in RPMI and thereafter infected with *C. auris* at a MOI 50 by sedimenting the yeast cells at 200 xg for 3 min and incubated at 37 °C, 5% CO_2_, and 90% humidity for 5, 10, 20, 40, 60, 80 min before fixation with 3.7 % PFA in PBS. *C. auris* cells were stained either with Calcofluor-white (5µM) or DAPI (1µg/ml).

### Antibodies and constructs

For details on antibodies and constructs, see Table S2.

### Inside-outside staining

Following infection, samples were fixed with methanol free PFA 4% for 10 min and washed three times with PBS. Incompletley internalized *C. auris* cells were stained with ConcanavalinA-568 (2.5 µg/ml in PBS) for 20 min at room temperature in the dark. Samples were washed three time with PBS and further stained with AlexaFluor488-phalloidin.

### Lysotracker assay

Cells were seeded on µ-Slide 8 wells (ibidi, Martinsried, Germany) at a density of 1x10^5^ in RPMI 1640 medium at 37 °C, 5% CO_2_, and 90% humidity and subsequently preincubated with 0.5 µM LysoTracker-Red DND-99 (Invitrogen, Massachusetts, USA) solution in RPMI with respective inhibitors or DMSO as control. Preconditioned cells were infected with *C.auris* prestained with Calcofluor-white (5 µM) in RPMI 1640 for 30 min. Optical Z-stacks with astep size of 0.3 µm were acquired from 20 min to 120 min post infection.

### Immunofluorescence and microscopy

Confluent macrophages were fixed after *C. auris* infection, and permeabilized for 5 min in 0.5% TritonX-100 in PBS. Thereafter cells were incubated at least for 30 min in blocking solution (5% normal human serum, 5% normal goat serum, 2% BSA in 0.05% Triton-X-100 in PBS), and incubated for at least 60 min at room temperature with the primary antibody in blocking solution. Cells were washed three times in 0.05% Triton-X-100 in PBS and then incubated for 60 min with the secondary antibody in blocking solution supplemented with AlexaFluor488/568/647-phalloidin (1:100), and optionally DAPI (1µg/ml) (Biotium, Fremont, USA) to label *C. auris* and cell nuclei. Alternatively. *C. auris* were prestained with Calcofluor-white (5µM) (Biotium, Fremont, USA) for 30 min. After three washes in PBS, coverslips were mounted on glass slides with FluoromountG (Invitrogen Massachusetts, USA). Images of fixed samples were acquired with a confocal laser-scanning microscope Olympus FV3000 equipped with an 60x UPlanApo HR Oil objective and Olympus FV-3000 software, Visitron SD-TIRF equipped with 100x CFI Plan Apo Lambda and Visitron software, and Velocity software. Images were processed with Imaris 7.6.1 and 9.9.1 versions and Fiji software.

### Macrophage electroporation

After detachment with Accutase, cells were washed in DPBS and resuspended in R-Buffer (at a concentration of 1x10^6^ cells per 100 µl buffer, 10 µg DNA) contained in the Neon Transfection System (Invitrogen, Massachusetts, USA). Macrophages were transiently transfected with plasmid DNA with the following settings: voltage 1000 V; width 40 ms; and 2 pulses. Transfected cells were resuspended in RPMI and seeded on glass bottom 8-Well µ-slide, 1x10^5^ cells per well. The cells were left to adhere for 1 h at 37°C, 5% CO_2_, and 90% humidity. Thereafter, 0.3 ml of monocyte culture medium was added, and cells were incubated for at least 5 h up to overnight for overexpression of the respective plasmid (see Table S2).

### Live imaging microscopy

Cells infected with *C. auris* were imaged in RPMI 1640 medium at 37°C, 5% CO_2_, and 90% humidity. Image acquisition was performed with Visitron SD-TIRF. Z-stacks were acquired for 20 min, with 30 s intervals, z-step size: 0.3 µm, in respective fluorescent channels. Fiji Trackmate plugin was used for semiautomatic detection of phagocytic podosomes and their track duration in z-stacks over time. Particle diameter was set at 0.5 µm, frame to frame linking was set at 1 µm with identical maximal gap-closing distance and maximal frame gap of 2 frames.

### Statistical Evaluation

Statistical evaluation of datasets was performed using a two-tailed Student’s *t* test or one-way ANOVA in GraphPad Prism 10.2.3 (GraphPad Software).

## Acknowledgements

We thank Frank Bentzien (UKE transfusion medicine) for buffy coats, Andrea Mordhorst and Kevin Schiemang for expert technical assistance, the UKE microscopy facility (umif) for help with microscopy and image analysis, Sergio Grinstein and Spencer Freeman for hosting K.S. for an internship at SickKids (Toronto, CAN), and Martin Aepfelbacher for continuous support. This work is part of the doctoral theses of K.S., R.B, and Q.S., and was supported by Deutsche Forschungsgemeinschaft (RTG2771/P4).

The authors declare no competing financial interests.

## Author Contributions

K.S., R.B., Q.S., and P.C designed and performed experiments, S.L. and P.C. designed experiments, K.S. prepared figures, and S.L. wrote the manuscript.

## Supplementary Figure Legends

**Suppl. Figure S1. Uptake of *C. albicans* is associated with the formation of phagocytic podosomes.** (A-D) Confocal micrographs of primary human macrophages infected with *C. albicans* cells, stained for F-actin using Alexa488-phalloidin (A,B), and stained for DNA with DAPI (C), with respective merge (D). Whiate box in (A) indicates detail regions shown in (B-D). Scale bars: 10 µm for (A), 2 µm for detail images

**Suppl. Figure S2. Hyphal outgrowth of *C. albicans* within primary macrophages.** Confocal micrographs (projection of optical z-stacks) of primary macrophages, fixed at indicated time points post infection with cells of *C. albicans* (A-I) or with *C. auris* (J-R), and stained for F-actin, after infection, with yeast cell wall stained with Calcofluor (F). Time points after infection are indicated on the left. Note hyphal growth of internalized *C. albicans,* but not of *C. auris,* with subsequent outgrowth from macrophages and establishment of an extensive fungal network. Scale bar: 50 µm.

**Suppl. Figure S3. Further components of *C. auris*-associated phagocytic podosomes.** (A-G) Confocal images of *C. auris-*containing phagosomes, with specimens stained for F-actin using Alexa488-phalloidin and for indicated components using specific antibodies. White boxes in overview images indicate detail regions shown on the right, with individual channels in black/white, and colored merges. Graphs show radial and tangential line scans of respective fluorescence intensities at individual F-actin foci. From top to bottom: cortactin (core; A), talin-1 (ring; B), zyxin (cap; C), LSP1 (cap; D), WASp (base; E), (phosphotyrosine; regulatory signal, F,G) with phospho-tyrosine visualized in confocal mode (F) or by STED superresolution microscopy (G). Scale bars: 2 µm for overviews and 1 µm for detail images See also Fig. 2.

**Suppl. Figure** S4. Test of inhibitors for disruption of ventral podosomes. Confocal images of primary macrophages stained for F-actin (A-F) or phosphotyrosine (G-L), with merges (M-R), treated with cytochalasin D, to inhibit actin dynamics (B,H,N), with CK666 (C,I,O) or CK869 (DJ,P), inhibitors of Arp2/3 complex, with wiskostatin, an inhibitor of WASp (E,K,O), or with PP2, an inhibitor of Src tyrosine kinases (F,L,P); scale bars: 50 µm. (S-V) Respective graphs show numbers of ventral podosomes per 1x10^3^ µm^2^. N≥ 3x 10^4^ µm^2^, from each time 3 donors. Values are given as mean ± SEM; *P<0.05; **P<0.01, according to Wilcoxon test. For specific values, see Suppl. Table 1. (see also Fig. 4).

## Supplementary Videos

**Suppl. Video S1. 3D animation of *Candida auris*-containing phagosome with associated phagocytic podosomes.** Detail images from primary human macrophage infected with *C. auris* cells, and stained for F-actin to visualize podosomes (green), and DAPI to visualize yeast DNA (blue), (See also Fig. 1I-K) Scale bar: 1 µm

**Suppl. Video S2. Formation and turnover of phagocytic podosomes during uptake of *Candida auris*.** Live cell video of primary human macrophage expressing lifeact-GFP to label F-actin, which is enriched at phagocytic podosomes, and infected with multiple *C. auris* cells. Maximum projection of optical z-stack with substraction of adhesive plane containing ventral podosomes. Experiment duration: 20 min, frame rate: 1/30 sec. (See also Fig. 3C-F).

## References

1 Meis, J. F. & Chowdhary, A. Candida auris: a global fungal public health threat. Lancet Infect Dis 18, 1298–1299, doi:10.1016/S1473-3099(18)30609-1 (2018).

2 Satoh, K. et al. Candida auris sp. nov., a novel ascomycetous yeast isolated from the external ear canal of an inpatient in a Japanese hospital. Microbiol Immunol 53, 41–44, doi:10.1111/j.1348-0421.2008.00083.x (2009).

3 Services, C. D. o. H. a. H. Antibiotic Resistance Threats in the United States (2019).

4 Eyre, D. W. et al. A Candida auris Outbreak and Its Control in an Intensive Care Setting. The New England journal of medicine 379, 1322–1331, doi:10.1056/NEJMoa1714373 (2018).

5 Di Lorenzo, A. et al. Candida auris cluster in a large third level Italian hospital: a case series. IJID Reg 13, 100468, doi:10.1016/j.ijregi.2024.100468 (2024).

6 Lockhart, S. R. et al. Simultaneous Emergence of Multidrug-Resistant Candida auris on 3 Continents Confirmed by Whole-Genome Sequencing and Epidemiological Analyses. Clin Infect Dis 64, 134–140, doi:10.1093/cid/ciw691 (2017).

7 Santana, D. J. & O’Meara, T. R. Forward and reverse genetic dissection of morphogenesis identifies filament-competent Candida auris strains. Nature communications 12, 7197, doi:10.1038/s41467-021-27545-5 (2021).

8 Huang, X. et al. Murine model of colonization with fungal pathogen Candida auris to explore skin tropism, host risk factors and therapeutic strategies. Cell host & microbe 29, 210–221 e216, doi:10.1016/j.chom.2020.12.002 (2021).

9 Horton, M. V., Holt, A. M. & Nett, J. E. Mechanisms of pathogenicity for the emerging fungus Candida auris. PLoS pathogens 19, e1011843, doi:10.1371/journal.ppat.1011843 (2023).

10 Santana, D. J., et al. A Candida auris-specific adhesin, Scf1, governs surface association, colonization, and virulence. Science 381, 1461–1467, doi:10.1126/science.adf8972 (2023).

11 Horton, M. V. et al. Candida auris Cell Wall Mannosylation Contributes to Neutrophil Evasion through Pathways Divergent from Candida albicans and Candida glabrata. mSphere 6, e0040621, doi:10.1128/mSphere.00406-21 (2021).

12 Johnson, C. J., Davis, J. M., Huttenlocher, A., Kernien, J. F. & Nett, J. E. Emerging Fungal Pathogen Candida auris Evades Neutrophil Attack. mBio 9, doi:10.1128/mBio.01403-18 (2018).

13 Uribe-Querol, E. & Rosales, C. Phagocytosis: Our Current Understanding of a Universal Biological Process. Frontiers in immunology 11, 1066, doi:10.3389/fimmu.2020.01066 (2020).

14 Fountain, A., Inpanathan, S., Alves, P., Verdawala, M. B. & Botelho, R. J. Phagosome maturation in macrophages: Eat, digest, adapt, and repeat. Adv Biol Regul 82, 100832, doi:10.1016/j.jbior.2021.100832 (2021).

15 Jaumouille, V. & Grinstein, S. Molecular Mechanisms of Phagosome Formation. Microbiol Spectr 4, doi:10.1128/microbiolspec.MCHD-0013-2015 (2016).

16 Allen, L. A. & Aderem, A. Molecular definition of distinct cytoskeletal structures involved in complement- and Fc receptor-mediated phagocytosis in macrophages. J Exp Med 184, 627–637, doi:10.1084/jem.184.2.627 (1996).

17 Linder, S. & Barcelona, B. Get a grip: Podosomes as potential players in phagocytosis. European journal of cell biology 102, 151356, doi:10.1016/j.ejcb.2023.151356 (2023).

18 Ostrowski, P. P., Freeman, S. A., Fairn, G. & Grinstein, S. Dynamic Podosome-Like Structures in Nascent Phagosomes Are Coordinated by Phosphoinositides. Developmental cell 50, 397–410 e393, doi:10.1016/j.devcel.2019.05.028 (2019).

19 Labrousse, A. M. et al. Frustrated phagocytosis on micro-patterned immune complexes to characterize lysosome movements in live macrophages. Frontiers in immunology 2, 51, doi:10.3389/fimmu.2011.00051 (2011).

20 Tertrais, M. et al. Phagocytosis is coupled to the formation of phagosome-associated podosomes and a transient disruption of podosomes in human macrophages. European journal of cell biology 100, 151161, doi:10.1016/j.ejcb.2021.151161 (2021).

21 Vorselen, D. et al. Phagocytic ‘teeth’ and myosin-II ‘jaw’ power target constriction during phagocytosis. eLife 10, doi:10.7554/eLife.68627 (2021).

22 Herron, J. C. et al. Actin nano-architecture of phagocytic podosomes. Nature communications 13, 4363, doi:10.1038/s41467-022-32038-0 (2022).

23 Roncero, C. & Duran, A. Effect of Calcofluor white and Congo red on fungal cell wall morphogenesis: in vivo activation of chitin polymerization. J Bacteriol 163, 1180–1185, doi:10.1128/jb.163.3.1180-1185.1985 (1985).

24 Linder, S., Cervero, P., Eddy, R. & Condeelis, J. Mechanisms and roles of podosomes and invadopodia. Nature reviews. Molecular cell biology 24, 86–106, doi:10.1038/s41580-022-00530-6 (2023).

25 Cervero, P. et al. Myosins 1e/f at the podosome base regulate podosome dynamics and promote macrophage migration. European journal of cell biology 104, 151514, doi:10.1016/j.ejcb.2025.151514 (2025).

26 Linder, S. et al. The polarization defect of Wiskott-Aldrich syndrome macrophages is linked to dislocalization of the Arp2/3 complex. Journal of immunology 165, 221–225 (2000).

27 Tehrani, S., Faccio, R., Chandrasekar, I., Ross, F. P. & Cooper, J. A. Cortactin has an essential and specific role in osteoclast actin assembly. Molecular biology of the cell 17, 2882–2895, doi:10.1091/mbc.e06-03-0187 (2006).

28 Marchisio, P. C. et al. Vinculin, talin, and integrins are localized at specific adhesion sites of malignant B lymphocytes. Blood 72, 830–833 (1988).

29 Gaidano, G. et al. Integrin distribution and cytoskeleton organization in normal and malignant monocytes. Leukemia 4, 682–687 (1990).

30 del Rio, A. et al. Stretching single talin rod molecules activates vinculin binding. Science 323, 638–641, doi:10.1126/science.1162912 (2009).

31 van den Dries, K., et al. Modular actin nano-architecture enables podosome protrusion and mechanosensing. Nature communications 10, 5171, doi:10.1038/s41467-019-13123-3 (2019).

32 Joosten, B., Willemse, M., Fransen, J., Cambi, A. & van den Dries, K. Super-Resolution Correlative Light and Electron Microscopy (SR-CLEM) Reveals Novel Ultrastructural Insights Into Dendritic Cell Podosomes. Frontiers in immunology 9, 1908, doi:10.3389/fimmu.2018.01908 (2018).

33 Cervero, P., Wiesner, C., Bouissou, A., Poincloux, R. & Linder, S. Lymphocyte-specific protein 1 regulates mechanosensory oscillation of podosomes and actin isoform-based actomyosin symmetry breaking. Nature communications 9, 515, doi:10.1038/s41467-018-02904-x (2018).

34 Linder, S., Nelson, D., Weiss, M. & Aepfelbacher, M. Wiskott-Aldrich syndrome protein regulates podosomes in primary human macrophages. Proceedings of the National Academy of Sciences of the United States of America 96, 9648–9653 (1999).

35 Wiesner, C., Faix, J., Himmel, M., Bentzien, F. & Linder, S. KIF5B and KIF3A/KIF3B kinesins drive MT1-MMP surface exposure, CD44 shedding, and extracellular matrix degradation in primary macrophages. Blood 116, 1559–1569, doi:10.1182/blood-2009-12-257089 (2010).

36 Pal, K., Zhao, Y., Wang, Y. & Wang, X. Ubiquitous membrane-bound DNase activity in podosomes and invadopodia. The Journal of cell biology 220, doi:10.1083/jcb.202008079 (2021).

37 Bhuwania, R. et al. Supervillin couples myosin-dependent contractility to podosomes and enables their turnover. Journal of cell science 125, 2300–2314, doi:10.1242/jcs.100032 (2012).

38 Gavazzi, I., Nermut, M. V. & Marchisio, P. C. Ultrastructure and gold-immunolabelling of cell-substratum adhesions (podosomes) in RSV-transformed BHK cells. Journal of cell science 94 (Pt 1), 85–99 (1989).

39 Linder, S. & Aepfelbacher, M. Podosomes: adhesion hot-spots of invasive cells. Trends Cell Biol 13, 376–385 (2003).

40 Weber, K., Hey, S., Cervero, P. & Linder, S. The circle of life: Phases of podosome formation, turnover and reemergence. European journal of cell biology 101, 151218, doi:10.1016/j.ejcb.2022.151218 (2022).

41 Jaumouille, V. & Waterman, C. M. Physical Constraints and Forces Involved in Phagocytosis. Frontiers in immunology 11, 1097, doi:10.3389/fimmu.2020.01097 (2020).

42 Mousa, G. Y., Trevithick, J. R., Bechberger, J. & Blair, D. G. Cytochalasin D induces the capping of both leukaemia viral proteins and actin in infected cells. Nature 274, 808–809, doi:10.1038/274808a0 (1978).

43 Yang, Q., Zhang, X. F., Pollard, T. D. & Forscher, P. Arp2/3 complex-dependent actin networks constrain myosin II function in driving retrograde actin flow. The Journal of cell biology 197, 939–956, doi:10.1083/jcb.201111052 (2012).

44 Cao, L., Huang, S., Basant, A., Mladenov, M. & Way, M. CK-666 and CK-869 differentially inhibit Arp2/3 iso-complexes. EMBO Rep 25, 3221–3239, doi:10.1038/s44319-024-00201-x (2024).

45 Leung, D. W., Morgan, D. M. & Rosen, M. K. Biochemical properties and inhibitors of (N-)WASP. Methods in enzymology 406, 281–296, doi:10.1016/S0076-6879(06)06021-6 (2006).

46 Linder, S., Hufner, K., Wintergerst, U. & Aepfelbacher, M. Microtubule-dependent formation of podosomal adhesion structures in primary human macrophages. Journal of cell science 113 Pt 23, 4165–4176 (2000).

47 Korade-Mirnics, Z. & Corey, S. J. Src kinase-mediated signaling in leukocytes. Journal of leukocyte biology 68, 603–613 (2000).

48 Cervero, P., Panzer, L. & Linder, S. Podosome reformation in macrophages: assays and analysis. Methods in molecular biology 1046, 97–121, doi:10.1007/978-1-62703-538-5_6 (2013).

49 Tkacz, J. S., Cybulska, E. B. & Lampen, J. O. Specific staining of wall mannan in yeast cells with fluorescein-conjugated concanavalin A. J Bacteriol 105, 1–5, doi:10.1128/jb.105.1.1-5.1971 (1971).

50 Westman, J., Moran, G., Mogavero, S., Hube, B. & Grinstein, S. Candida albicans Hyphal Expansion Causes Phagosomal Membrane Damage and Luminal Alkalinization. mBio 9, doi:10.1128/mBio.01226-18 (2018).

51 Weerasinghe, H. et al. Candida auris uses metabolic strategies to escape and kill macrophages while avoiding robust activation of the NLRP3 inflammasome response. Cell reports 42, 112522, doi:10.1016/j.celrep.2023.112522 (2023).

52 Cervero, P. et al. Myo1e/f at the podosome base regulate podosome dynamics and promote macrophage migration. bioRxiv, 2025.2004.2028.651090, doi:10.1101/2025.04.28.651090 (2025).

53 Proag, A., Bouissou, A., Vieu, C., Maridonneau-Parini, I. & Poincloux, R. Evaluation of the force and spatial dynamics of macrophage podosomes by multi-particle tracking. Methods 94, 75–84, doi:10.1016/j.ymeth.2015.09.002 (2016).

54 Le, P. H. et al. Surface Architecture Influences the Rigidity of Candida albicans Cells. Nanomaterials (Basel) 12, doi:10.3390/nano12030567 (2022).

55 Sridharan, R., Cavanagh, B., Cameron, A. R., Kelly, D. J. & O’Brien, F. J. Material stiffness influences the polarization state, function and migration mode of macrophages. Acta Biomater 89, 47–59, doi:10.1016/j.actbio.2019.02.048 (2019).

56 Maxson, M. E. et al. Integrin-based diffusion barrier separates membrane domains enabling the formation of microbiostatic frustrated phagosomes. eLife 7, doi:10.7554/eLife.34798 (2018).

57 Depierre, M., Pompili, C. & Niedergang, F. Phagocytosis at a glance. Journal of cell science 138, doi:10.1242/jcs.263833 (2025).

58 Lukacsi, S. et al. The differential role of CR3 (CD11b/CD18) and CR4 (CD11c/CD18) in the adherence, migration and podosome formation of human macrophages and dendritic cells under inflammatory conditions. PloS one 15, e0232432, doi:10.1371/journal.pone.0232432 (2020).

59 Soloviev, D. A. et al. Identification of pH-regulated antigen 1 released from Candida albicans as the major ligand for leukocyte integrin alphaMbeta2. Journal of immunology 178, 2038–2046, doi:10.4049/jimmunol.178.4.2038 (2007).

60 Diamond, R. D. Interactions of phagocytic cells with Candida and other opportunistic fungi. Arch Med Res 24, 361–369 (1993).

61 Gomez-Gaviria, M., Martinez-Alvarez, J. A., Chavez-Santiago, J. O. & Mora-Montes, H. M. Candida haemulonii Complex and Candida auris: Biology, Virulence Factors, Immune Response, and Multidrug Resistance. Infect Drug Resist 16, 1455–1470, doi:10.2147/IDR.S402754 (2023).

62 Tsuboi, S. & Meerloo, J. Wiskott-Aldrich syndrome protein is a key regulator of the phagocytic cup formation in macrophages. The Journal of biological chemistry 282, 34194–34203, doi:10.1074/jbc.M705999200 (2007).

63 van den Dries, K., Linder, S., Maridonneau-Parini, I. & Poincloux, R. Probing the mechanical landscape - new insights into podosome architecture and mechanics. Journal of cell science 132, doi:10.1242/jcs.236828 (2019).

64 van den Dries, K., et al. Geometry sensing by dendritic cells dictates spatial organization and PGE(2)-induced dissolution of podosomes. Cellular and molecular life sciences: CMLS 69, 1889–1901, doi:10.1007/s00018-011-0908-y (2012).

65 Linder, S. & Wiesner, C. Tools of the trade: podosomes as multipurpose organelles of monocytic cells. Cellular and molecular life sciences: CMLS, doi:10.1007/s00018-014-1731-z (2014).

66 Jaumouille, V., Cartagena-Rivera, A. X. & Waterman, C. M. Coupling of beta2 integrins to actin by a mechanosensitive molecular clutch drives complement receptor-mediated phagocytosis. Nat Cell Biol 21, 1357–1369, doi:10.1038/s41556-019-0414-2 (2019).

67 Kopp, P. et al. The kinesin KIF1C and microtubule plus ends regulate podosome dynamics in macrophages. Molecular biology of the cell 17, 2811–2823, doi:10.1091/mbc.E05-11-1010 (2006).

68 El Azzouzi, K., Wiesner, C. & Linder, S. Metalloproteinase MT1-MMP islets act as memory devices for podosome reemergence. The Journal of cell biology 213, 109–125, doi:10.1083/jcb.201510043 (2016).

69 Majeed, M., Caveggion, E., Lowell, C. A. & Berton, G. Role of Src kinases and Syk in Fcgamma receptor-mediated phagocytosis and phagosome-lysosome fusion. Journal of leukocyte biology 70, 801–811 (2001).

