## Supplementary figures and images for "Phagocytic podosomes enable efficient uptake of *Candida auris* by primary human macrophages"

### Suppl. Figures 1-4

#### Suppl. Figure 1

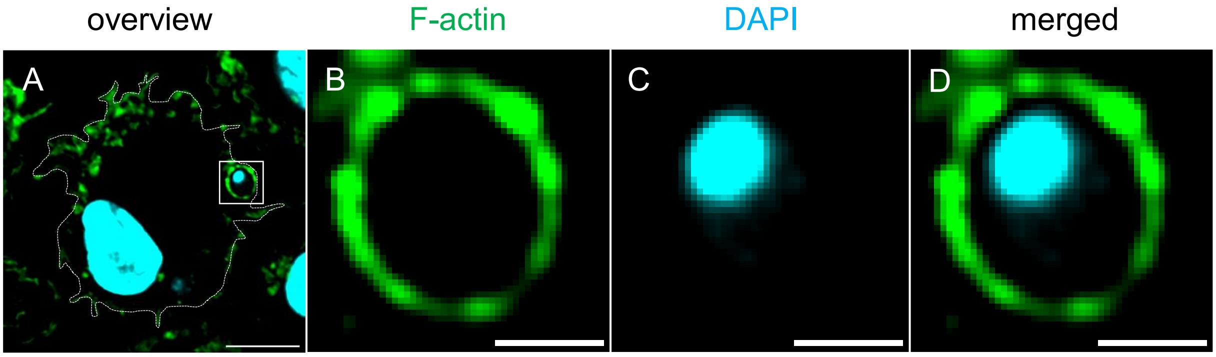

Suppl. Figure 2

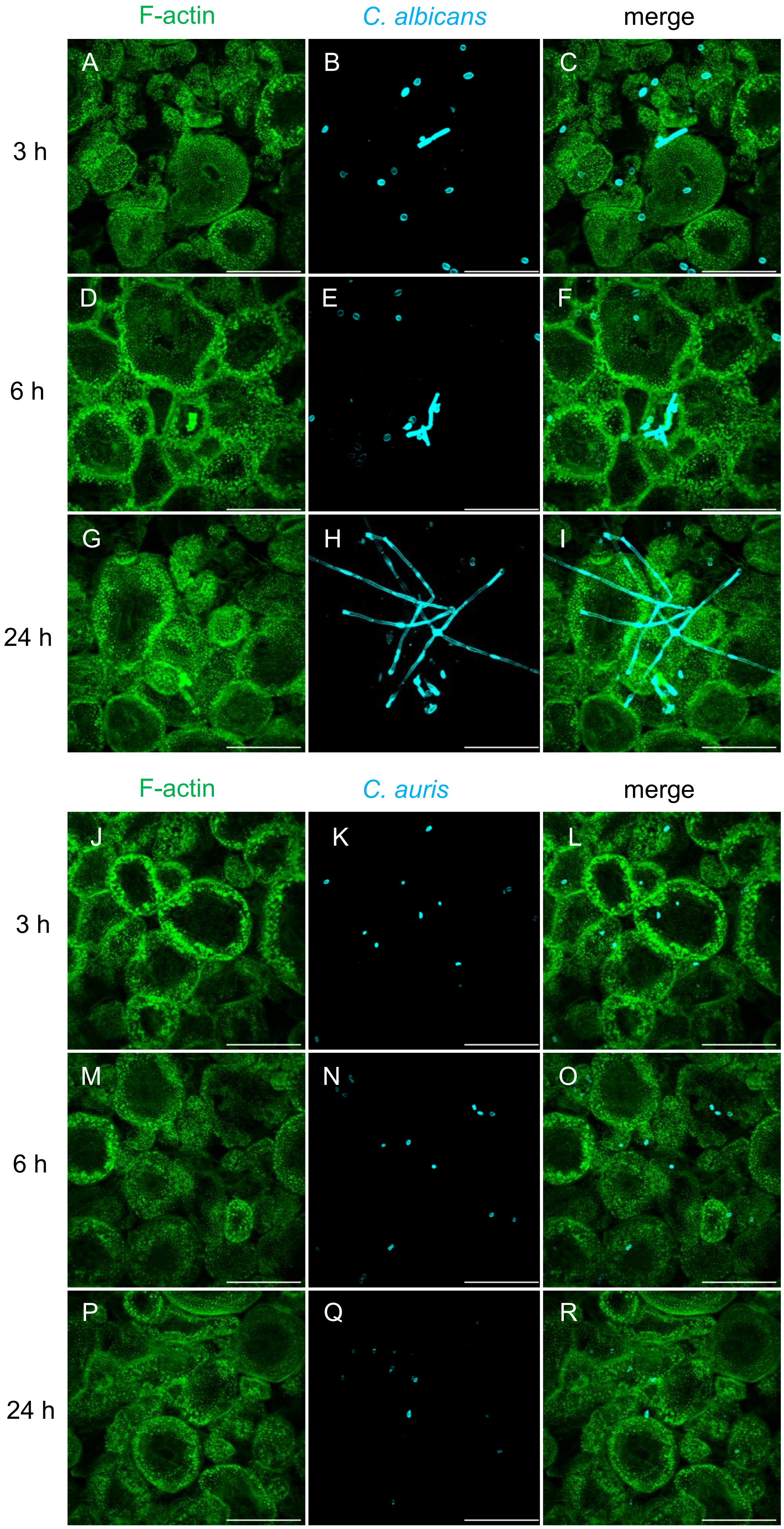

#### Suppl. Figure 3

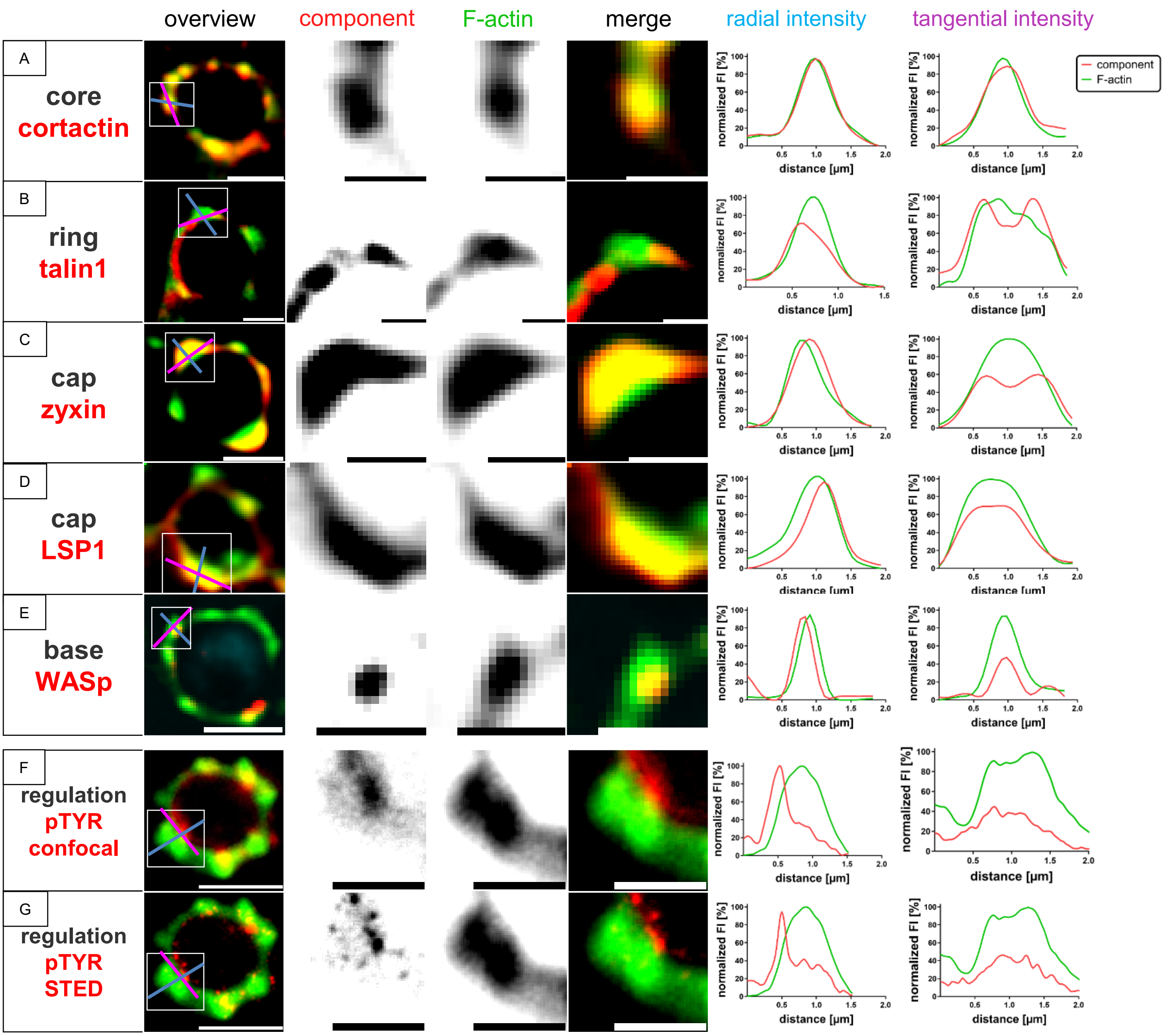

#### Suppl. Figure 4

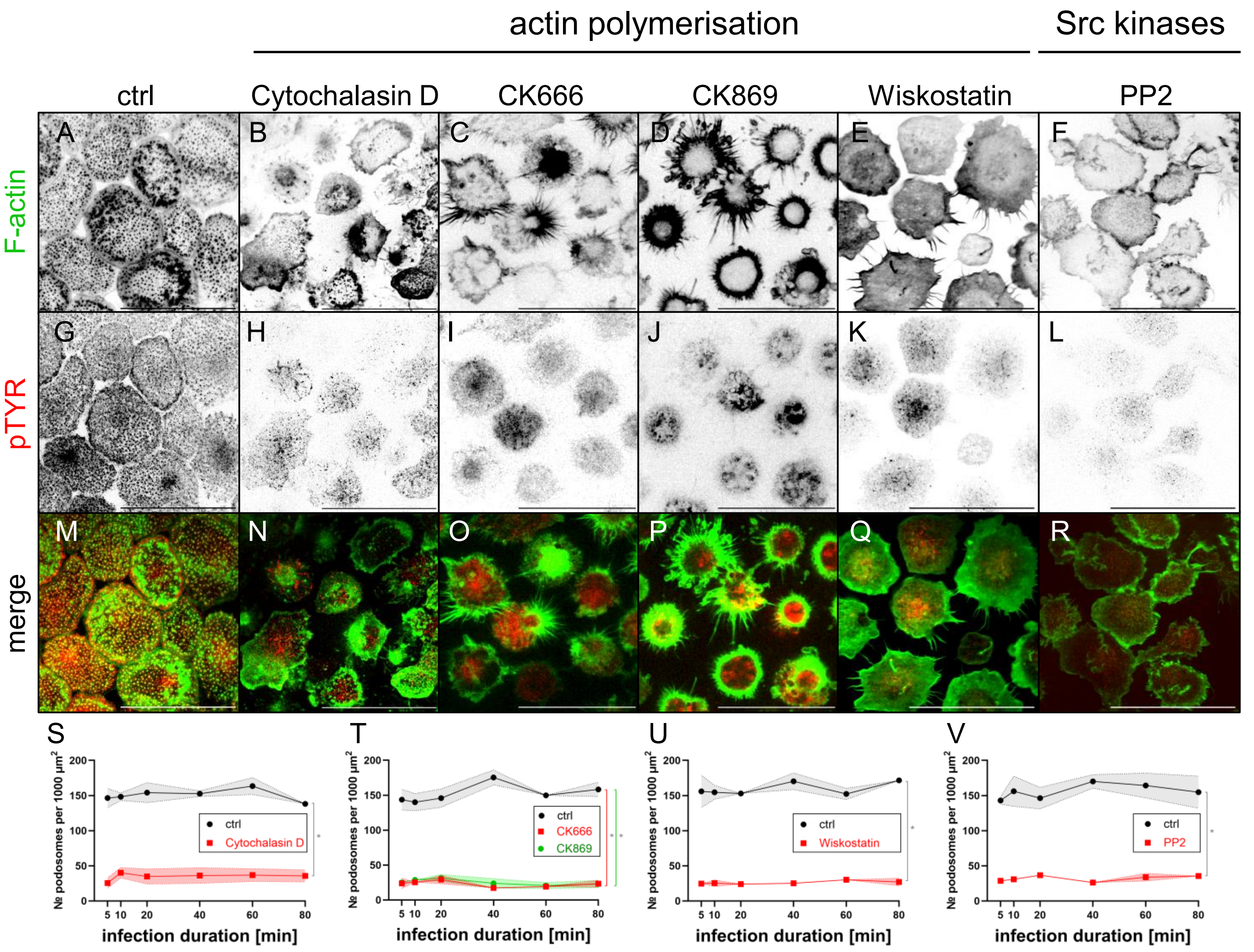
