## Supplementary material for "Phagocytic podosomes enable efficient uptake of *Candida auris* by primary human macrophages": Suppl. Tables 1+2

| **Figure 3 A.** *C. auris* phagosomes with phagocytic podosomes. (Mean ±S.E.M.) | | | | | | | | | | | | | | |
| --- | --- | --- | --- | --- | --- | --- | --- | --- | --- | --- | --- | --- | --- | --- |
| Infection duration [min] | | | Percentage of *C. auris* with phagocytic podosomes [%] | | |  | | | | | | | | |
| 5 | | | 66.37±4.56 | | |  | | | | | | | | |
| 10 | | | 75.27±3.89 | | |  | | | | | | | | |
| 20 | | | 65.40±8.70 | | |  | | | | | | | | |
| 40 | | | 26.03±9.21 | | |  | | | | | | | | |
| 60 | | | 4.90±4.00 | | |  | | | | | | | | |
| **Figure 3B.** № of phagocytic podosomes. (Mean ±S.E.M. [%]) | | | | | | | | | | | | | | |
| Infection duration [min] | | | Mean № of phagocytic podosomes per *Candida* phagosome | | |  | | | | | | | | |
| 5 | | | 3.12±0.28 | | |  | | | | | | | | |
| 10 | | | 3.28±0.21 | | |  | | | | | | | | |
| 20 | | | 2.94±0.30 | | |  | | | | | | | | |
| 40 | | | 1.12±0.20 | | |  | | | | | | | | |
| 60 | | | 0.16±0.08 | | |  | | | | | | | | |
| **Figure 3F.** Lifetime of phagocytic podosomes | | | | | | | | | | | | | | |
| Track duration [min] | | | | Frequency distribution [%] | | | | | Track duration [min] | | | | Frequency distribution [%] | |
| 1 | | | | 30.77 | | | | | 11 | | | | 1.01 | |
| 2 | | | | 19.87 | | | | | 12 | | | | 0.64 | |
| 3 | | | | 15.75 | | | | | 13 | | | | 0.73 | |
| 4 | | | | 10.44 | | | | | 14 | | | | 0.09 | |
| 5 | | | | 7.69 | | | | | 15 | | | | 0.00 | |
| 6 | | | | 4.58 | | | | | 16 | | | | 0.09 | |
| 7 | | | | 2.75 | | | | | 17 | | | | 0.09 | |
| 8 | | | | 2.38 | | | | | 18 | | | | 0.09 | |
| 9 | | | | 1.65 | | | | | 19 | | | | 0.27 | |
| 10 | | | | 0.92 | | | | | 20 | | | | 0.18 | |
| **Figure 4 A-D.** *Candida* phagosomes with phagocytic podosomes. (Mean ±S.E.M. [%]) | | | | | | | | | | | | | | |
| Infection duration [min] | ctrl | | | Cytochalasin D [1.5µM] | ctrl | | | CK666  [100 µM] | | CK869  [100 µM] | ctrl | Wiskostatin [50 µM] | ctrl | PP2  [25 µM] |
| 5 | 71.99 ± 4.64 | | | 0.00±0.00 | 47.13±1.75 | | | 6.94±3.00 | | 4.76±3.89 | 57.00±8.41 | 0.00±0.00 | 66.48±9.25 | 5.56±2.62 |
| 10 | 52.18 ±0.53 | | | 0.00±0.00 | 55.06±6.17 | | | 12.74±4.51 | | 0.00±0.00 | 50.26±5.99 | 0.00±0.00 | 50.33±1.73 | 12.91±4.61 |
| 20 | 22.49±4.81 | | | 5.56±4.54 | 23.13±2.68 | | | 9.48±6.39 | | 8.47±3.54 | 22.64±3.04 | 0.00±0.00 | 27.74±4.59 | 14.64±3.93 |
| 40 | 6.05±1.82 | | | 11.62±4.76 | 14.34±2.08 | | | 8.15±3.47 | | 12.75±7.37 | 10.04±1.28 | 0.00±0.00 | 16.19±1.71 | 11.12±1.55 |
| 60 | 7.19±0.62 | | | 12.50±5.89 | 7.37±0.99 | | | 8.83±5.56 | | 15.00±6.24 | 4.45±2.02 | 0.00±0.00 | 8.10±2.74 | 10.71±2.02 |
| 80 | 1.05±0.18 | | | 10.32±4.25 | 2.76±1.43 | | | 5.97±4.38 | | 8.07±3.34 | 1.21±0.50 | 0.00±0.00 | 0.72±0.59 | 4.87±1.43 |
| **Figure 4 E-H.** № of phagocytic podosomes per phagosome. (Mean ±S.E.M.) | | | | | | | | | | | | | | |
| Infection duration [min] | ctrl | | | Cytochalasin D [1.5µM] | ctrl | | | CK666 [100 µM] | | CK869 [100 µM] | ctrl | Wiskostatin [50 µM] | ctrl | PP2  [25 µM] |
| 5 | 2.83±0.57 | | | 0.00±0.00 | 2.15±0.35 | | | 0.10±0.04 | | 0.10±0.08 | 2.37±0.40 | 0.00±0.00 | 2.57±0.32 | 0.15±0.08 |
| 10 | 1.98±0.15 | | | 0.00±0.00 | 2.49±0.32 | | | 0.16±0.02 | | 0.06±0.05 | 1.94±0.35 | 0.00±0.00 | 1.82±0.09 | 0.17±0.06 |
| 20 | 0.99±0.23 | | | 0.06±0.05 | 1.01±0.14 | | | 0.15±0.11 | | 0.22±0.10 | 0.98±0.11 | 0.00±0.00 | 1.12±0.13 | 0.42±0.12 |
| 40 | 0.38±0.07 | | | 0.34±0.16 | 0.76±0.12 | | | 0.17±0.07 | | 0.28±0.14 | 0.34±0.07 | 0.00±0.00 | 0.73±0.09 | 0.24±0.01 |
| 60 | 0.49±0.10 | | | 0.19±0.11 | 0.38±0.06 | | | 0.37±0.10 | | 0.55±0.23 | 0.10±0.04 | 0.00±0.00 | 0.33±0.05 | 0.23±0.04 |
| 80 | 0.07±0.02 | | | 0.15±0.07 | 0.36±0.11 | | | 0.11±0.07 | | 0.20±0.08 | 0.04±0.02 | 0.00±0.00 | 0.04±0.04 | 0.09±0.03 |
| **Figure 4 I-L.** Internalization of *C. auris* upon use of inhibitors. (Mean ±S.E.M. [%]) | | | | | | | | | | | | | | |
| Infection duration [min] | ctrl | | | Cytochalasin D [1.5µM] | ctrl | | | CK666  [100 µM] | | CK869  [100 µM] | ctrl | Wiskostatin [50 µM] | ctrl | PP2  [25 µM] |
| 5 | 33.93±7.72 | | | 4.56±1.95 | 30.59±5.78 | | | 14.71±3.12 | | 6.87±1.32 | 36.83±2.51 | 2.02±1.65 | 36.50±5.15 | 18.25±4.60 |
| 10 | 41.41±4.42 | | | 1.99±0.83 | 37.76±9.18 | | | 15.30±3.17 | | 5.32±1.25 | 43.81±4.72 | 1.38±0.69 | 48.39±3.04 | 19.34±3.07 |
| 20 | 68.54±6.04 | | | 2.82±0.90 | 49.61±3.19 | | | 23.50±4.03 | | 9.53±0.56 | 67.19±3.51 | 2.59±0.43 | 69.10±4.60 | 37.94±4.77 |
| 40 | 77.75±6.53 | | | 7.70±1.91 | 68.85±3.49 | | | 18.21±4.61 | | 16.44±1.47 | 82.48±4.25 | 4.22±1.47 | 90.87±4.14 | 54.37±0.63 |
| 60 | 86.82±6.37 | | | 13.38±1.03 | 75.49±2.74 | | | 20.90±1.21 | | 14.59±2.62 | 88.18±2.66 | 5.06±2.56 | 91.28±4.23 | 68.47±3.07 |
| 80 | 92.10±4.53 | | | 9.03±1.30 | 84.83±3.82 | | | 28.69±3.54 | | 28.65±3.24 | 85.53±3.22 | 4.58±0.97 | 95.19±2.83 | 69.07±3.52 |
| **Figure 4 M-P.** Phagocytic index upon use of inhibitors. (Mean ±S.E.M. [%]) | | | | | | | | | | | | | | |
| Infection duration [min] | ctrl | | | Cytochalasin D [1.5µM] | ctrl | | | CK666  [100 µM] | | CK869  [100 µM] | ctrl | Wiskostatin [50 µM] | ctrl | PP2  [25 µM] |
| 5 | 154.44±41.73 | | | 12.22±4.80 | 150.00±49.91 | | | 41.11±5.52 | | 20.00±1.57 | 120.00±6.85 | 4.44±3.63 | 183.33±16.63 | 45.56±7.09 |
| 10 | 162.22±23.53 | | | 5.56±2.40 | 156.67±46.85 | | | 67.78±27.32 | | 18.89±0.91 | 123.33±17.00 | 3.33±1.57 | 202.22±17.31 | 63.33±13.97 |
| 20 | 277.78±39.51 | | | 10.00±4.16 | 303.33±51.09 | | | 90.00±4.16 | | 36.67±4.16 | 232.22±17.31 | 5.56±0.91 | 261.11±26.17 | 146.67±38.91 |
| 40 | 344.45±21.33 | | | 22.22±6.35 | 381.11±58.63 | | | 96.67±42.28 | | 83.33±21.60 | 327.78±20.75 | 10.00±3.14 | 390.00±22.17 | 190.00±23.47 |
| 60 | 525.56±12.70 | | | 37.78±11.90 | 523.33±64.94 | | | 95.56±25.06 | | 66.66±27.22 | 508.89±50.81 | 14.44±7.75 | 471.11±35.43 | 217.78±19.65 |
| 80 | 531.11±31.75 | | | 25.55±3.27 | 493.33±59.71 | | | 141.11±53.20 | | 126.67±20.43 | 494.44±37.89 | 13.34±2.72 | 544.44±53.46 | 244.44±35.36 |
| **Suppl. Figure 4 S-V.** Number of ventral podosomes per 1000 µm^2^ upon use of inhibitors. (Mean ±S.E.M. [%]) | | | | | | | | | | | | | | |
| Infection duration [min] | ctrl | | | Cytochalasin D [1.5µM] | ctrl | | | CK666  [100 µM] | | CK869  [100 µM] | ctrl | Wiskostatin [50 µM] | ctrl | PP2  [25 µM] |
| 5 | 146.51±11.13 | | | 25.60±6.05 | 143.68±12.16 | | | 23.90±6.24 | | 25.18±3.17 | 155.94±18.96 | 24.68±0.21 | 142.99±4.37 | 28.75±2.47 |
| 10 | 148.40±4.41 | | | 40.25±6.20 | 139.94±10.06 | | | 25.39±2.60 | | 28.86±3.00 | 154.63±8.10 | 25.35±3.84 | 156.21±17.57 | 30.87±2.22 |
| 20 | 154.23±11.65 | | | 34.87±8.89 | 145.83±10.76 | | | 29.23±5.00 | | 31.47±4.08 | 153.02±2.37 | 24.11±1.79 | 146.33±12.45 | 36.72±1.07 |
| 40 | 152.69±3.24 | | | 36.26±9.22 | 175.43±8.73 | | | 17.19±3.19 | | 23.95±5.68 | 170.27±9.65 | 25.22±2.18 | 170.20±8.00 | 26.24±1.42 |
| 60 | 163.50±9.86 | | | 36.88±7.56 | 149.82±2.36 | | | 19.21±2.16 | | 20.65±3.59 | 152.28±6.51 | 30.26±2.64 | 164.25±14.46 | 33.94±4.66 |
| 80 | 138.21±0.68 | | | 35.65±7.09 | 158.29±8.48 | | | 23.46±4.08 | | 22.85±4.42 | 171.60±1.01 | 27.02±4.43 | 154.86±18.61 | 35.55±3.11 |
| **Figure 5U**. *C.auris*- F-actin distances. (Mean ±S.E.M [µm]) | | | | | | | | | | | | |  | |
| ctrl | | | | CK666 [100µM] | | | | PP2 [25µM] | | | EDTA [1mM] | |  | |
| 0.27±0.02 | | | | 0.53±0.03 | | | | 0.45±0.02 | | | 0.045±0.03 | |  |  |
| **Figure 5 E.J.O.T.** Cumulative intensity of *C.auris*- F-actin distances. (Mean ±S.E.M. [%]) | | | | | | | | | | | | | | |
| Distance from center [µm] | | ctrl  F-actin | | ctrl  *C.auris* cell wall | CK666 [100µM]  F-actin | | CK666  [100µM]  *C.auris* cell wall | | | PP2  [25 µM]  F-actin | PP2  [25 µM] *C.auris* cell wall | EDTA  [1mM]  F-actin | EDTA [1mM] *C.auris* cell wall |  |
| -1.21 | | 5.15±0.48 | | 22.38±1.06 | 5.58±0.74 | | 25.60±1.34 | | | 5.73±0.68 | 26.00±1.88 | 6.34±0.77 | 25.98±1.22 |  |
| -1.15 | | 5.14±0.48 | | 22.78±1.07 | 5.58±0.75 | | 26.75±1.43 | | | 5.83±0.69 | 26.69±1.83 | 6.07±0.64 | 26.35±1.24 |  |
| -1.08 | | 5.55±0.67 | | 22.97±1.02 | 6.05±0.73 | | 27.65±1.41 | | | 6.59±0.73 | 27.49±1.85 | 6.89±0.76 | 26.84±1.24 |  |
| -1.02 | | 5.67±0.58 | | 23.83±1.06 | 5.76±0.66 | | 29.08±1.49 | | | 6.96±0.76 | 28.02±1.82 | 6.99±0.73 | 27.59±1.23 |  |
| -0.96 | | 6.30±0.59 | | 24.16±1.02 | 6.43±0.86 | | 30.94±1.51 | | | 6.94±0.78 | 29.72±1.91 | 7.90±0.81 | 28.94±1.26 |  |
| -0.89 | | 6.84±0.60 | | 25.61±1.06 | 6.78±0.92 | | 32.93±1.61 | | | 7.76±0.82 | 30.92±1.99 | 9.21±1.02 | 30.04±1.32 |  |
| -0.83 | | 7.45±0.67 | | 26.96±1.13 | 7.59±0.95 | | 35.28±1.74 | | | 8.41±0.86 | 31.91±2.00 | 10.04±1.02 | 31.91±1.35 |  |
| -0.77 | | 8.36±0.76 | | 28.19±1.12 | 7.78±0.95 | | 39.40±1.92 | | | 9.04±0.88 | 34.90±2.08 | 10.00±0.92 | 33.84±1.49 |  |
| -0.7 | | 8.13±0.61 | | 30.46±1.15 | 8.23±0.93 | | 43.64±2.20 | | | 9.79±0.87 | 38.60±2.23 | 10.30±0.87 | 36.60±1.61 |  |
| -0.64 | | 9.29±0.79 | | 32.81±1.23 | 8.78±1.04 | | 49.55±2.50 | | | 10.51±0.87 | 42.82±2.34 | 12.26±1.12 | 41.04±1.81 |  |
| -0.57 | | 9.87±0.72 | | 37.04±1.37 | 9.43±0.96 | | 55.91±2.66 | | | 11.52±0.92 | 47.74±2.49 | 12.79±0.91 | 46.29±1.92 |  |
| -0.51 | | 11.50±0.76 | | 42.03±1.46 | 10.33±0.99 | | 62.65±2.71 | | | 13.04±1.03 | 54.36±2.50 | 14.50±0.97 | 52.19±1.99 |  |
| -0.45 | | 14.12±1.06 | | 47.16±1.56 | 11.27±0.99 | | 70.20±2.51 | | | 14.32±1.07 | 62.54±2.54 | 16.07±1.12 | 58.87±2.07 |  |
| -0.38 | | 16.23±1.13 | | 55.63±1.71 | 13.17±1.08 | | 77.83±2.29 | | | 16.47±1.09 | 70.56±2.36 | 18.19±1.27 | 67.91±2.20 |  |
| -0.32 | | 19.99±1.37 | | 65.74±1.81 | 14.75±1.12 | | 83.72±2.25 | | | 18.87±1.27 | 79.88±2.06 | 21.33±1.42 | 77.44±2.21 |  |
| -0.26 | | 24.62±1.50 | | 77.32±1.86 | 17.58±1.19 | | 86.29±2.48 | | | 22.33±1.36 | 87.93±1.83 | 24.44±1.36 | 84.77±2.22 |  |
| -0.19 | | 32.29±1.60 | | 87.84±1.67 | 21.58±1.35 | | 83.01±2.89 | | | 27.63±1.66 | 89.50±2.09 | 29.17±1.58 | 87.56±2.46 |  |
| -0.13 | | 43.46±1.82 | | 93.58±1.36 | 27.16±1.68 | | 73.61±3.27 | | | 34.09±1.92 | 83.38±2.68 | 35.85±1.88 | 83.87±2.98 |  |
| -0.06 | | 58.71±2.11 | | 90.45±1.60 | 35.94±2.24 | | 59.61±3.48 | | | 43.25±2.26 | 70.89±3.28 | 46.22±2.24 | 73.60±3.41 |  |
| 0 | | 74.78±2.15 | | 78.93±2.31 | 47.40±2.67 | | 44.50±3.13 | | | 55.51±2.46 | 55.85±3.49 | 58.45±2.47 | 58.66±3.47 |  |
| 0.06 | | 88.99±1.63 | | 61.81±2.62 | 61.20±2.92 | | 31.46±2.48 | | | 68.45±2.30 | 41.85±3.28 | 71.39±2.47 | 42.86±2.99 |  |
| 0.13 | | 94.66±1.11 | | 44.49±2.39 | 73.16±2.78 | | 21.50±1.73 | | | 82.33±1.86 | 30.54±2.87 | 84.48±1.93 | 29.20±2.26 |  |
| 0.19 | | 90.38±1.45 | | 29.53±2.07 | 82.00±2.37 | | 14.96±1.16 | | | 92.91±1.36 | 21.42±2.34 | 89.84±1.54 | 19.24±1.59 |  |
| 0.26 | | 80.04±2.29 | | 19.93±1.78 | 86.21±2.12 | | 10.62±0.82 | | | 93.27±1.19 | 15.77±1.95 | 89.41±1.53 | 12.85±1.23 |  |
| 0.32 | | 66.05±2.95 | | 14.19±1.27 | 85.46±2.14 | | 7.80±0.56 | | | 84.98±1.74 | 12.29±1.71 | 82.47±2.29 | 9.44±0.91 |  |
| 0.38 | | 53.15±3.16 | | 10.07±0.80 | 79.79±2.62 | | 6.10±0.42 | | | 73.00±2.48 | 9.26±1.26 | 74.02±2.81 | 6.58±0.62 |  |
| 0.45 | | 40.98±2.98 | | 7.57±0.52 | 70.97±3.07 | | 5.05±0.33 | | | 59.79±2.96 | 7.57±1.18 | 65.24±3.19 | 5.07±0.45 |  |
| 0.51 | | 31.45±2.59 | | 5.71±0.43 | 62.72±3.32 | | 4.35±0.33 | | | 48.29±3.17 | 6.44±1.07 | 56.21±3.15 | 4.28±0.44 |  |
| 0.57 | | 24.89±2.35 | | 5.18±0.45 | 54.99±3.59 | | 3.76±0.27 | | | 40.23±3.03 | 5.30±0.89 | 49.69±3.10 | 3.71±0.37 |  |
| 0.64 | | 21.00±2.24 | | 4.54±0.43 | 47.77±3.57 | | 3.32±0.22 | | | 33.70±2.86 | 4.51±0.68 | 44.09±3.01 | 3.07±0.34 |  |
| 0.7 | | 17.96±2.11 | | 3.88±0.45 | 40.67±3.29 | | 2.87±0.19 | | | 29.45±2.81 | 4.05±0.52 | 42.12±3.11 | 2.69±0.31 |  |
| 0.77 | | 15.38±1.95 | | 3.87±0.40 | 35.98±3.12 | | 2.48±0.22 | | | 26.80±2.75 | 3.42±0.43 | 35.90±2.74 | 2.70±0.32 |  |
| 0.83 | | 13.97±2.09 | | 2.98±0.27 | 32.61±3.03 | | 2.33±0.22 | | | 25.51±2.77 | 3.25±0.43 | 35.17±2.73 | 2.26±0.27 |  |
| 0.89 | | 13.27±2.02 | | 3.18±0.42 | 29.22±2.90 | | 2.10±0.19 | | | 23.98±2.73 | 3.04±0.38 | 31.83±2.68 | 2.33±0.28 |  |
| 0.96 | | 12.00±1.83 | | 3.16±0.52 | 25.94±2.59 | | 2.04±0.19 | | | 23.17±2.77 | 2.54±0.32 | 29.26±2.65 | 2.36±0.27 |  |
| 1.02 | | 10.94±1.84 | | 3.06±0.59 | 24.19±2.39 | | 1.97±0.20 | | | 22.57±2.75 | 2.27±0.35 | 28.76±2.72 | 2.09±0.30 |  |
| 1.08 | | 10.89±1.93 | | 2.95±0.67 | 22.30±2.29 | | 1.89±0.19 | | | 21.84±2.71 | 2.19±0.26 | 25.84±2.50 | 1.98±0.32 |  |
| 1.15 | | 10.66±1.64 | | 2.48±0.65 | 21.31±2.22 | | 1.74±0.20 | | | 22.52±2.73 | 1.86±0.32 | 23.44±2.32 | 2.22±0.34 |  |
| 1.21 | | 9.82±1.51 | | 2.47±0.68 | 20.41±2.07 | | 1.76±0.19 | | | 22.20±2.69 | 1.78±0.32 | 23.29±2.33 | 2.23±0.44 |  |
| **Figure 6 M.N.** LAMP1 enrichment at phagocytic podosomes upon use of inhibitors. (Mean ±S.E.M[%]) | | | | | | | | | |  | **Figure 6 A2.B2.** Lysotracker intensity upon use of inhibitors. (Mean ±S.E.M [a.u.]) | | | |
| Infection duration [min] | ctrl | | | CK666  [100µM] | ctrl | | | PP2  [25 µM] | |  | ctrl | CK666 [100µM] | ctrl | PP2  [25µM] |
| 5 | 38.99±3.84 | | | 10.72±4.46 | 39.30±14.30 | | | 5.26±4.30 | |  | 396.48±11.09 | 208.98±7.22 | 342.69±11.10 | 187.92±7.94 |
| 10 | 50.90±7.14 | | | 15.48±6.80 | 53.23±12.54 | | | 19.76±6.30 | |  |  |  |  |  |
| 20 | 57.78±4.10 | | | 33.40±6.82 | 62.32±7.02 | | | 33.39±7.65 | |  |  |  |  |  |
| 40 | 72.61±5.78 | | | 35.36±6.64 | 82.63±5.81 | | | 45.17±15.33 | |  |  |  |  |  |
| 60 | 82.22±5.08 | | | 53.11±9.02 | 92.93±1.78 | | | 58.86±17.45 | |  |  |  |  |  |
| 80 | 89.36±3.45 | | | 60.08±5.11 | 93.38±1.12 | | | 62.47±10.37 | |  |  |  |  |  |

| **Suppl. Table 2. Antibodies and Constructs** | | | | |
| --- | --- | --- | --- | --- |
| Antibodies and staining reagents | | | | |
| Antibody target | Company | Product number | Species | Dilution for Immunoluorescence |
| arp2 | Abcam | ab49674 | mouse | 1:400 |
| vinculin | Sigma-Aldrich | V9264 | mouse | 1:100 |
| CD11b/Mac-1 | BD Pharmingen | 550374 | mouse | 1:400 |
| a-actinin | Santa Cruz | sc-17829 | mouse | 1:100 |
| myosin If (B-5) | Santa Cruz | sc-376534 | mouse | 1:100 (methanole prefixation required) |
| MT1-MMP (MMP-14) | Merck Millipore | MAB3328 | mouse | 1:400 |
| DnaseX (DNASE1L1) | Abnova | H0001774-M02 | mouse | 1:100 |
| myosinIIA | Sigma-Aldrich | M8064 | rabbit | 1:100 |
| pTyr | Santa Cruz | sc-7020 | mouse | 1:200 |
| cortactin | BD Transduction Laboratories | 610049 | mouse | 1:100 |
| talin-1 | Bio-RAD | 97H6 | mouse | 1:100 |
| zyxin | Thermo Scientific | 39-6000 | mouse | 1:100 |
| LSP1 | AtlasAntibodies (Sigma) | HPA019693 | rabbit | 1:100 |
| WASp | Santa Cruz | sc-13139 | mouse | 1:100 (methanole prefixation required) |
| Anti-mouse-Abberior STAR RED | Abberior | STRED | goat | 1:100 |
| Anti-rabbit-AlexaFluor 488/568/647 | Thermo Scientific |  | donkey or goat | 1:200 |
| Anti-mouse-AlexaFluor 488/568/647 | Thermo Scientific |  | donkey or goat | 1:200 |
| Phalloidin AlexaFluor488/568/647 | Thermo Scientific |  |  | 1:100-1:400 |
| Calcofluor | Biotium |  |  | 5 µM |
| DAPI | Biotium |  |  | 1 µg/ml |
| Plasmids | | | | |
| Plasmid | Provided by |  |  |  |
| Lifeact-GFP | M.Sixt |  |  |  |
